# glmmDMR reveals replicate-level methylation variance as a major determinant of false-positive DMR detection

**DOI:** 10.64898/2026.06.29.734667

**Authors:** Yuki Daito, Mako Uechi, Tetsu Kinoshita, Kaoru Tonosaki

## Abstract

**Background:** Accurate identification of differentially methylated regions (DMRs) is fundamental to epigenomic research but remains challenging due to biological variability among replicates, heterogeneous effect sizes, and the tendency of adjacent cytosines to share similar methylation states. Many existing methods aggregate methylation measurements before statistical testing or do not explicitly account for replicate-level variability, contributing to elevated false-positive rates.

**Results:** We developed glmmDMR, a DMR detection framework that combines generalized linear mixed models with a seed-based strategy for reconstructing DMRs from locally high-confidence signals while explicitly modeling replicate-level variability. Using simulated datasets with known ground-truth DMRs, we demonstrate that false-positive detections are more strongly associated with methylation variance among biological replicates than with the magnitude of methylation differences between groups. glmmDMR achieved higher precision than existing approaches while maintaining competitive recall, particularly for subtle methylation differences. Site-level modeling with beta regression provided the strongest overall performance, and seed-based region construction reduced artificial DMR fragmentation, improving recovery of true DMR boundaries and producing more contiguous, biologically interpretable DMRs. Applied to *Arabidopsis thaliana ddm1* methylomes and a rice DEMETER-LIKE DNA demethylase mutant (*Osdml3a-1*), glmmDMR identified biologically meaningful DMRs, revealing widespread TE-associated hypomethylation and subtle TE-family-specific hypermethylation.

**Conclusions:** Replicate-level methylation variance is an important determinant of DMR detection performance, and explicitly modeling this variance improves discrimination of biologically meaningful methylation changes from high-variance signals. By combining variance-aware statistical modeling with seed-based region construction, glmmDMR provides a robust framework for identifying contiguous, biologically interpretable DMRs across diverse methylome datasets.

## Background

DNA methylation is a conserved epigenetic modification that regulates gene expression, transposable element (TE) silencing, and developmental programs across eukaryotes [1]. In plants, DNA methylation occurs in CG-, CHG-, and CHH-sequence contexts, which differ substantially in their maintenance mechanisms and genomic distributions [1, 2]. The identification of differentially methylated regions (DMRs) across biological conditions is a central goal of epigenomic research, with applications ranging from the characterization of natural epigenetic variation and developmental transitions to the functional annotation of epigenetic mutants [3, 4]. However, extracting biological meaning from differences in methylation levels between distinct samples remains challenging because the relationship between DNA methylation and downstream biological outcomes is often complex and context-dependent [5, 6], and spurious or poorly defined DMRs may confound efforts to link epigenetic variation to biological outcomes. Improving the reliability of DMR detection is therefore a prerequisite for more interpretable downstream analyses.

Despite its importance, accurate DMR detection remains statistically challenging. Genome-wide profiling of DNA methylation by whole-genome bisulfite sequencing (WGBS) and related technologies [7, 8] has enabled the large-scale identification of DMRs across diverse biological contexts. However, methylation data are characterized by greater-than-expected variability (overdispersion), biological variability among replicates, heterogeneous size effects across genomic regions, and a complex spatial structure in which adjacent cytosines often exhibit correlated methylation states [9, 10]. These properties complicate statistical modeling and may contribute to instability in DMR calls, particularly across genomic regions that differ in the consistency of methylation levels among biological replicates Many computational tools have been developed to address these limitations. Site-level approaches that are widely used because of their simplicity and minimal distributional assumptions and operate on individual cytosines, such as methylKit [11] and Fisher’s exact test [8, 12], perform statistical testing at individual cytosines or within genomic windows, followed by region merging. However, these approaches are sensitive to low-coverage sites and do not explicitly model replicate-level variability. Region-level methods such as DSS [13, 14] improve statistical modeling through beta-binomial models that account for extra biological variability together with spatial smoothing but do not explicitly incorporate replicate effects within a mixed-effects framework. DMRfinder [15] identifies DMRs using local significance thresholds and region-merging procedures, whereas MACAU2 [16] incorporates mixed-model inference to account for overdispersion and sample structure. Systematic evaluations have shown that the performance of these methods varies substantially with effect size, sequencing depth, and dataset characteristics [17, 18]. A common challenge across many approaches is that statistical confidence is often determined primarily by effect size or aggregated significance measures, while the contribution of replicate-level variability is considered only indirectly during DMR prioritization and region construction. As a result, genomic regions with similar methylation differences between biological conditions may receive substantially different levels of statistical support depending on the consistency of methylation patterns among biological replicates.

A second challenge concerns the identification of DMR boundaries. Most existing methods define DMRs by merging neighboring cytosines or windows that are individually significant. Although effective in many situations, this strategy can be sensitive to local noise and may generate fragmented DMRs or inconsistent boundary placement [17]. Importantly, neighboring windows can differ substantially in statistical confidence despite showing similar methylation differences, particularly when replicate-level variability is high. Region construction strategies that initiate from locally high-confidence signals and constrain expansion based on directional and statistical consistency may therefore provide a more robust framework for capturing biologically meaningful methylation changes while limiting the propagation of noisy signals. However, such approaches remain relatively uncommon.

To address these challenges, we developed glmmDMR. This framework combines generalized linear mixed models (GLMMs), which explicitly model biological replicates as random effects, with a seed-based region-construction strategy that reconstructs biologically meaningful DMRs from locally high-confidence signals. By integrating variance-aware statistical modeling with biologically informed region construction, glmmDMR addresses both replicate-level methylation variance and artificial DMR fragmentation during DMR detection. Here, we benchmark glmmDMR against existing methods using simulated and real methylome datasets and show that combining variance-aware GLMM-based modeling with seed-based region construction improves the accuracy, robustness, and biological relevance of DMR detection.

## Results

### Overview of the glmmDMR framework for replicate-aware DMR detection

Accurate DMR detection is complicated by variation among biological replicates, which can substantially affect statistical support even for genomic regions with similar methylation differences between groups. In particular, neighboring genomic windows may receive markedly different statistical support despite exhibiting comparable methylation differences between groups, depending on the consistency of methylation patterns among biological replicates. To address this challenge, we developed glmmDMR, a framework that combines variance-aware statistical modeling with seed-based region construction (**Fig. 1**).

**Fig. 1.**
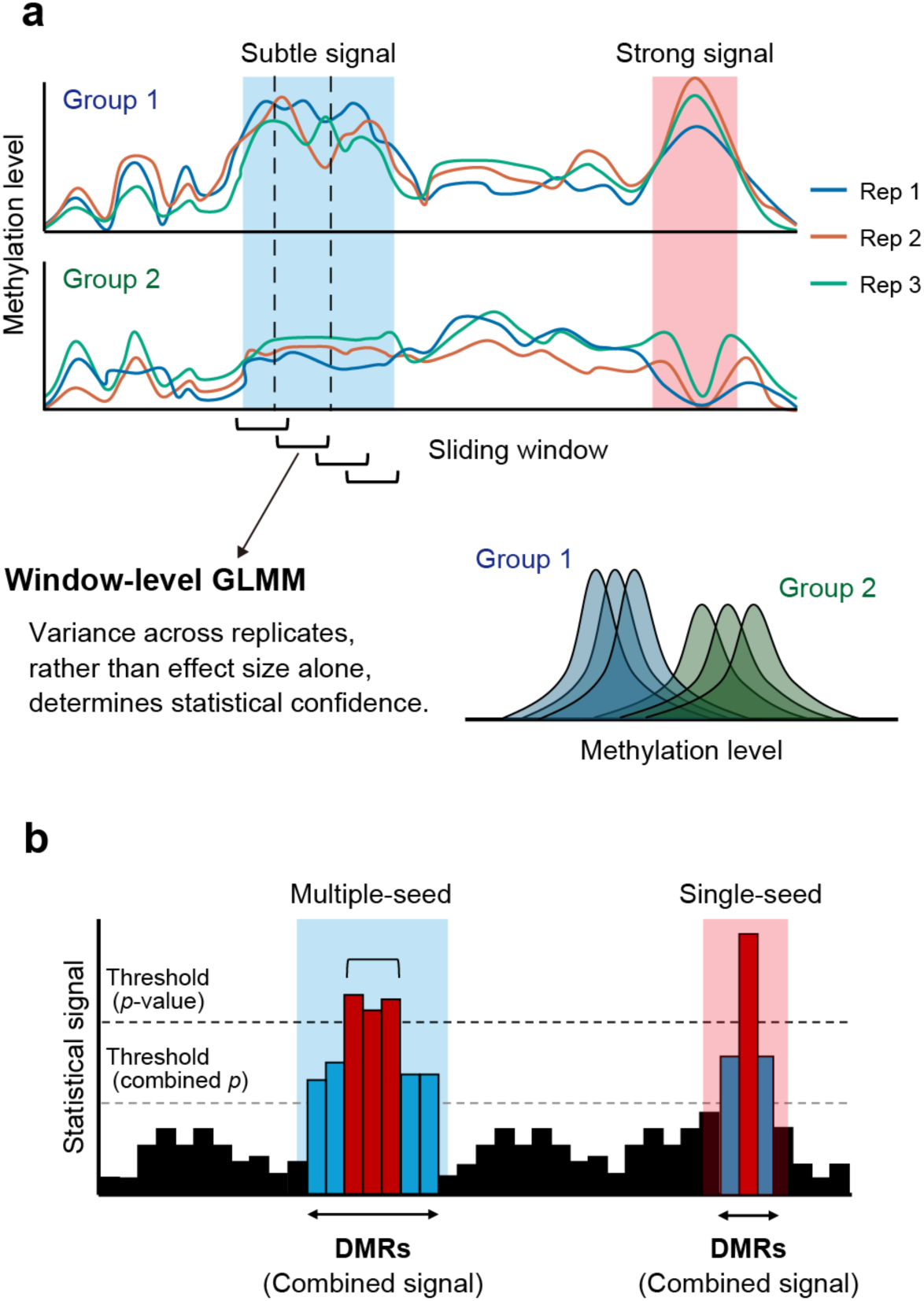
Conceptual overview of variance-aware DMR detection and seed-based region construction in glmmDMR. (**a**) Diagram illustrating the replicate-aware, window-level, methylation analysis. Methylation profiles from three biological replicates are shown for two groups (Group 1 and Group 2) across a representative genomic region. The highlighted regions indicate candidate DMRs subjected to window-level statistical testing using sliding windows applied across the genome. The blue-shaded region illustrates a methylation difference with low replicate-level variance, whereas the red-shaded region illustrates a similar methylation difference with higher replicate-level variance. Window-level statistical inference was performed using generalized linear mixed models (GLMMs), in which statistical confidence was determined by both the magnitude of the methylation difference and the replicate-level variance rather than by effect size alone. Distributions below the profiles show the replicate-level methylation distributions for Group 1 (blue) and Group 2 (green) at representative windows. (**b**) Conceptual illustration of seed-based DMR-construction strategies. Window-level statistical signals are integrated into DMRs using two complementary seed-based strategies. In the multiple-seed strategy, clusters of neighboring windows with strong statistical signals are jointly evaluated to define seed regions using combined *p*-values. In the single-seed strategy, individual highly significant windows are used as initial seeds. Starting from each seed, adjacent windows are then iteratively incorporated based on genomic proximity, consistency of DMR direction, and statistical support. Region-level statistical significance is evaluated using combined *p*-values throughout the expansion procedure. This framework enables the construction of biologically relevant DMRs while reducing the fragmentation associated with conventional window-merging strategies.

The first component of glmmDMR uses GLMMs [19] to evaluate methylation differences between the two experimental groups within sliding windows while explicitly modeling biological replicates as random effects (**Fig. 1a**). Under this framework, statistical confidence is determined by both the magnitude of methylation differences between groups and the variation among biological replicates. Consequently, regions that exhibit consistent methylation differences between groups across all replicates receive stronger statistical support than those with greater replicate-to-replicate variability.

The second component of glmmDMR merges significant windows into DMRs using a seed-based strategy (**Fig. 1b**). Rather than defining regions solely through threshold-based merging of neighboring windows [17], glmmDMR initiates DMR construction from locally high-confidence signals and iteratively expands regions by merging neighboring windows showing the same direction of methylation change (hyper- or hypomethylation), together with combined statistical support derived from the Stouffer method [20]. We implemented three complementary strategies: single-seed (SS), multiple-seed (MS), and a hybrid (HB) strategy that combines both approaches. These strategies enable the detection of both isolated high-confidence signals and broader regions with multiple locally significant windows while reducing DMR fragmentation and improving boundary definition compared with conventional merging approaches. glmmDMR supports multiple methylation sequencing platforms, including WGBS, enzymatic methyl sequencing (EM-seq) [21], and Oxford Nanopore Technologies (ONT) sequencing with methylation calls [22]. It provides four model configurations based on combinations of data representation (site level or aggregated) and response distribution (binomial or beta regression). Together, these components provide a unified framework for variance-aware DMR detection by combining explicit modeling of methylation variance among biological replicates with seed-based region construction.

### Site-level data representation and beta regression maximize the performance of DMR detection

To evaluate the influence of the different modeling strategies on DMR detection performance, we generated simulated CG methylation datasets that capture key characteristics of real methylome data, including overdispersion, variation among biological replicates, and site-level heterogeneity [9] (see Methods; **Fig. S1)**.

Simulations were restricted to the CG context because the objective was to compare statistical model performance rather than context-specific methylation biology. We compared the four GLMM model configurations defined above [19], assessing window-level performance against the predefined simulated DMRs (ground truth).

Across all benchmarking metrics, the beta-site configuration, which retains site-level methylation information and models methylation proportions using beta regression, consistently outperformed all other configurations. This advantage was most pronounced in precision–recall (PR) analysis: the beta-site model maintained consistently higher precision across a broad range of recall values, whereas the aggregated and binomial models showed a rapid loss of precision as recall rose (**Fig. 2a**). Receiver operating characteristic (ROC) analysis corroborated these findings: the beta-site model showed a more pronounced extension toward the upper-left corner of the curve compared to the other three configurations, which followed closely overlapping trajectories (**Fig. 2b**). Power–false discovery rate (FDR) analysis further confirmed that the beta-site model achieved higher recall across a wide range of FDR thresholds, whereas the beta-aggregate model exhibited notably lower sensitivity under stringent thresholds (**Fig. 2c**). Together, these results indicate that preserving site-level information substantially improves detection performance and that beta regression provides additional gains over binomial modeling when site-level representations are used.

**Fig. 2.**
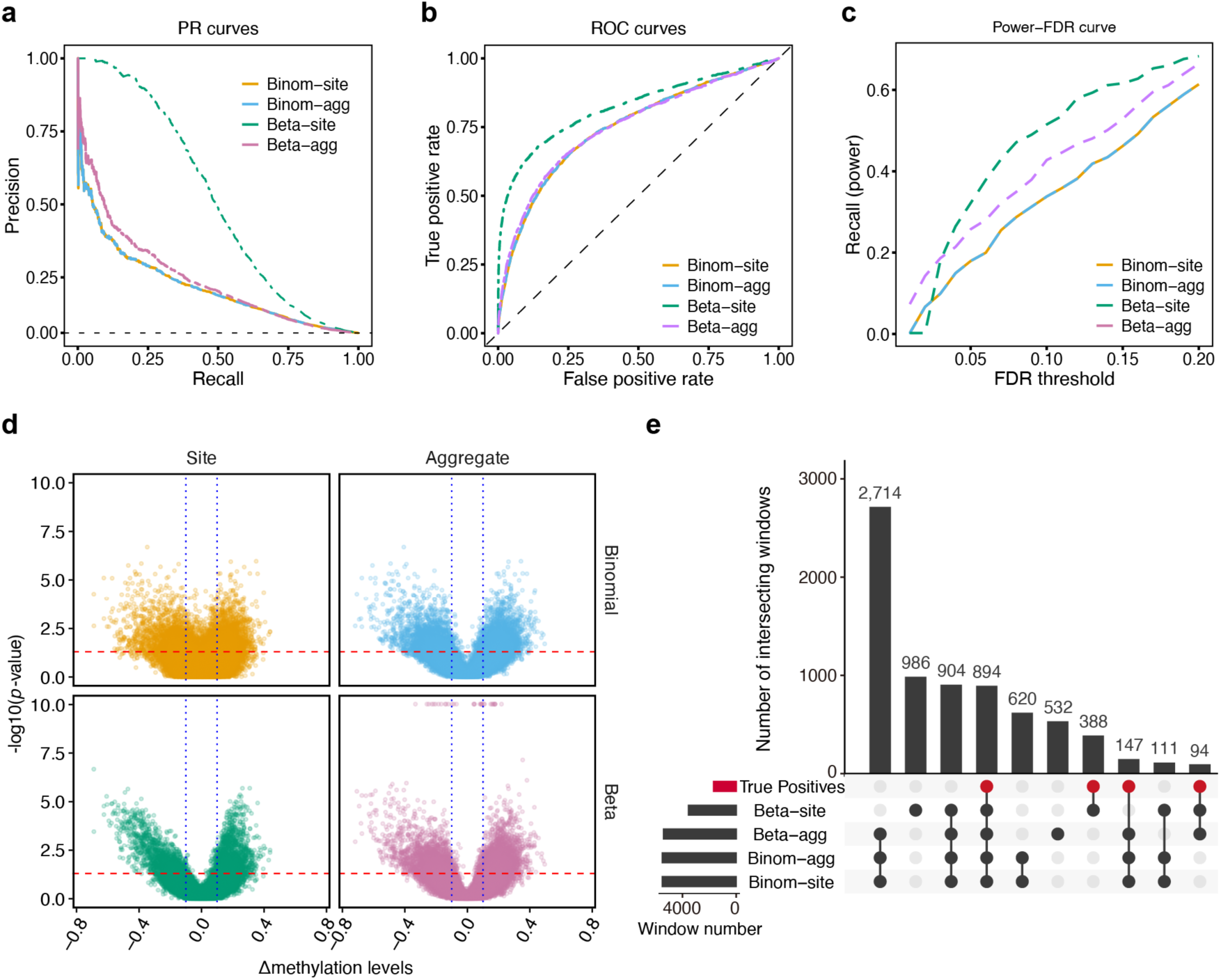
Performance of different GLMM model configurations using simulated methylome datasets. (**a**) Precision–recall (PR) curves for four GLMM model configurations evaluated using simulated CG methylome datasets. Curves were generated by varying the false discovery rate (FDR) threshold applied to window-level test statistics. Performance was compared among binomial and beta response distributions combined with site-level or aggregated window-level data representations (Binom-site, Binom-agg, Beta-site, and Beta-agg). The dashed horizontal line indicates the baseline precision, corresponding to the proportion of true-positive windows in the dataset. (**b**) Receiver operating characteristic (ROC) curves for the four GLMM model configurations. The dashed diagonal line indicates random classification performance. (**c**) Power–FDR relationship for each model configuration. Recall (power) was evaluated as the proportion of true-positive windows detected across higher FDR thresholds (0–0.20). (**d**) Relationship between methylation difference (Δmethylation) and window-level statistical significance (−log_10_[*p*-value]) for each GLMM configuration. Each point represents an individual genomic window. Red dashed horizontal lines indicate the significance threshold (*p* = 0.05); blue dashed vertical lines indicate |Δmethylation| = 0.1. Site-level and aggregated representations are shown separately for the binomial and beta models. (**e**) Overlap analysis of significant windows detected by the four GLMM configurations at FDR < 0.05. The UpSet plot shows the 10 largest intersection sets. Bar heights indicate the number of windows in each intersection. The red circles beneath the bars indicate the proportion of true-positive windows within each intersection set.

These differences in performance were not attributable to differences in global statistical calibration. Detection rates as a function of differences in methylation between groups (effect size) were broadly similar across configurations, with only modest differences at intermediate effect sizes (**Fig. S2a**). Similarly, the relationship between discovery counts and −log_10_(*p*) thresholds, as well as quantile–quantile (QQ) plot analyses, showed comparable behaviors across models, with only minor deviations (**Fig. S2b, c**). These observations indicate that the superior performance of the beta-site model is unlikely to be due to statistical inflation or calibration artifacts. We therefore investigated the characteristics of true-positive (TP) and false-positive (FP) detections to identify the factors underlying these differences in performance.

### False-positive detections are driven by methylation variance among replicates rather than absolute effect sizes

To identify the features that distinguish true from spurious detections, we compared methylation effect size, methylation variance, sequencing coverage, and coverage variance between TP and FP windows across all four GLMM configurations (**Fig. S3**). Although TP windows exhibited larger absolute methylation differences between groups than FP windows in all configurations (**Fig. S3a**), the separation between groups was moderate and incomplete, suggesting that effect size alone does not reliably distinguish genuine signals from spurious ones. In marked contrast, methylation variance among replicates was consistently significantly higher for FP windows than TP windows across all configurations (**Fig. S3b**). Neither sequencing coverage nor variance in coverage depth differed significantly between TP and FP windows (**Fig. S3c, d**), ruling out read depth as a confounding factor. These results indicate that methylation variance among replicates is a stronger discriminator between TP and FP windows than methylation effect size, identifying replicate-level variability as a major contributor to the detection of FP DMRs.

The strong association between elevated methylation variance and FP detection raised a question: if effect size is not the primary contributor to detection accuracy, why do model configurations differ in their performance? To address this question, we examined the extent of relationship between methylation effect size and statistical significance across model configurations (**Fig. 2d**). The beta-site model showed a clearer relationship between methylation effect size and statistical significance, consistent with its ability to prioritize windows with strong, consistent signals. By contrast, the aggregated models showed significant windows over a relatively narrower range of effect sizes. Notably, the beta-aggregate model exhibited a concentration of highly significant windows despite their relatively modest methylation differences between groups, suggesting lower discrimination between effect size and statistical significance under this configuration. These patterns are consistent with the superior precision observed with the beta-site model in PR analyses above and indicate that site-level data representation more effectively links statistical significance to the magnitude of true methylation differences between groups.

Overlap analysis revealed differences in the detection characteristics of the four configurations (**Fig. 2e**). A substantial number of detected windows were shared among all four configurations, indicating broad agreement in the detection of strong methylation signals. However, the beta-site model also recovered a subset of windows located within ground-truth DMRs that were not detected by the other configurations, consistent with its superior overall performance. By contrast, windows uniquely detected by the binomial and beta-aggregate models showed smaller overlaps with the shared core set of TP windows, indicating differences in detection behavior among model formulations.

Together, these analyses suggest that methylation variance among biological replicates is a major determinant of FP detection. The superior performance of the beta-site model appears to arise not from greater sensitivity to effect size alone but from its ability to more effectively align statistical significance with biologically meaningful methylation differences between groups, while reducing the influence of high-variance windows among replicates that are disproportionately associated with FP detections [9, 16]. glmmDMR addresses this issue by incorporating biological replicates as random effects within a GLMM framework, thereby providing a statistical mechanism for distinguishing consistent methylation changes from high-variance signals. In the following sections, we evaluate whether this variance-aware modeling improves region-level DMR detection and how seed-based integration further enhances DMR reconstruction.

### Seed-based integration strategies improve the accuracy of region-level DMR detection

Having established that methylation variance among replicates is a major contributor to FP detections at the window level, we asked whether seed-based region construction might improve region-level DMR detection by limiting the propagation of spurious window-level signals into reconstructed DMRs. We compared the three glmmDMR seed strategies (SS, MS, and HB) against conventional region-construction approaches based on *p*-value aggregation (Simes [23] and Stouffer [20]) and FDR-based window merging, using the same simulated datasets described above, for which the true DMR boundaries were known.

The methods differed substantially in both the numbers and structural properties of the reconstructed DMRs. Relative to the simulated ground-truth set (*n* = 435), conventional approaches generated either substantially fewer or substantially more DMRs than expected. FDR-based merging produced only 149 DMRs, whereas aggregation based on the Simes and Stouffer methods generated 1,061 and 1,592 DMRs, respectively (**Fig. 3a**). By contrast, the seed-based strategies within glmmDMR produced DMR counts much closer to the ground truth (SS, 420; MS, 309; HB, 401), indicating improved recovery of the expected regional structure. The length distribution of all defined DMRs further differentiated the methods. Seed-based approaches, particularly the MS and HB strategies, showed substantially smaller Kolmogorov–Smirnov distances (*D*) from the ground-truth distribution (MS, *D* = 0.12; HB, *D* = 0.12) than conventional methods (FDR, *D* = 0.73; Stouffer, *D* = 0.42; Simes, *D* = 0.30), suggestive of a more faithful reconstruction of true DMR sizes (**Fig. 3b**). Consistent with these observations, summary precision–recall performance showed that all three seed-based strategies achieved a more favorable balance between sensitivity and specificity than conventional approaches, with the MS strategy exhibiting the highest overall precision and overall F1 score among the seed-based strategies (**Fig. 3c**).

**Fig. 3.**
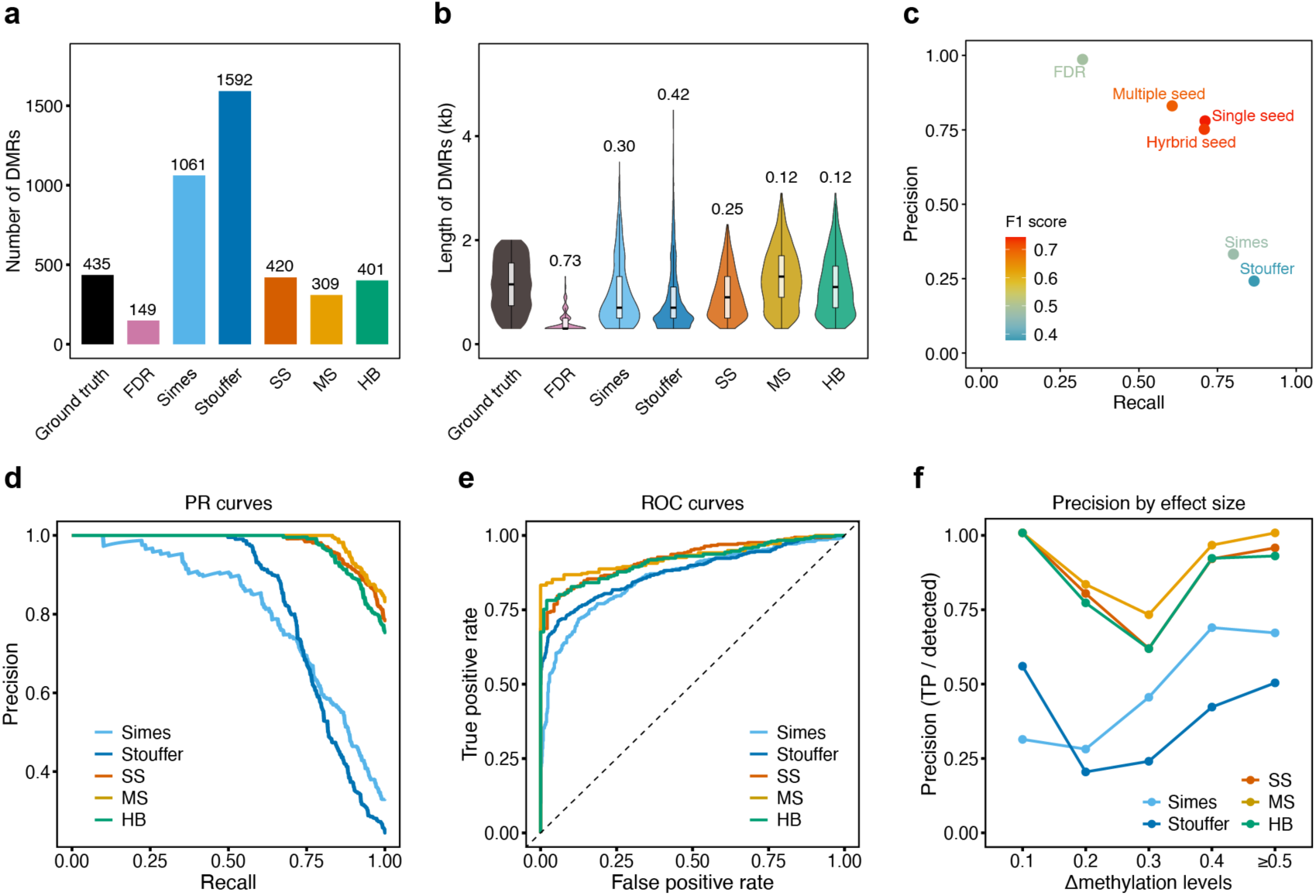
Performance comparison of seed-based DMR-construction strategies using simulated methylome datasets. (**a**) Numbers of DMRs detected by seed-based DMR-construction strategies (SS, single-seed; MS, multiple-seed; HB, hybrid) or conventional region-construction approaches (Simes, Stouffer, or FDR-based merging). The salmon bar indicates the number of ground-truth DMRs (*n* = 435). (**b**) Distribution of DMR length (kb) for each method and for the ground-truth DMRs. Violin plots show the full distribution; boxplots show the median (center line) and interquartile range (IQR; box), with whiskers extending to 1.5× IQR. *D* values shown above each violin plot indicate the Kolmogorov–Smirnov (KS) distance between the distribution of DMR length for each method and the ground-truth distribution. (**c**) Scatterplot showing the extent of correlation between recall and precision for each DMR-construction strategy at a fixed significance threshold. Point color indicates the F1 score (harmonic mean of precision and recall). (**d**) PR curves for each DMR-construction strategy. Curves were generated by varying the significance threshold applied to region-level combined *p*-values. (**e**) ROC curves for each DMR-construction strategy. The dashed diagonal line indicates random classification performance. (**f**) Precision as a function of methylation effect size (Δmethylation levels). Precision was calculated separately for detected DMRs grouped by the Δmethylation bin (0.1, 0.2, 0.3, 0.4, and ≥0.5) assigned to overlapping ground-truth DMRs.

Because FDR-based merging does not provide a region-level statistical score that can be varied continuously, we restricted subsequent threshold-dependent analyses to methods that generate region-level combined *p*-values. PR analysis demonstrated that all three seed-based strategies maintain substantially higher precision across a broad range of recall values. By contrast, the Simes and Stouffer methods exhibited rapid precision loss at higher recall levels (**Fig. 3d**). ROC analysis showed concordant trends, with the curves from seed-based strategies consistently above those from conventional methods (**Fig. 3e**). These differences were particularly pronounced at intermediate methylation effect sizes (Δ ≈ 0.1–0.3), where conventional approaches showed markedly lower precision whereas seed-based strategies maintained robust performance (**Fig. 3f**). This observation is especially relevant because many biologically meaningful methylation changes are not always associated with large methylation differences between groups.

The statistical basis for these improvements was supported by combined *p*-value analyses (**Fig. S4**). As significance thresholds rose, the number of discovered DMRs fell more gradually for seed-based approaches than conventional methods, consistent with more stable and statistically consistent region definitions. QQ plot analysis indicated a stronger enrichment of low combined *p*-values among DMRs generated from seed-based strategies, reflecting a greater concentration of true signals within their reconstructed regions.

FP DMRs exhibited greater methylation variance among replicates than TP DMRs across all strategies, consistent with the window-level analyses (**Fig. S5**). The higher precision achieved by seed-based strategies is therefore consistent with a lower incorporation of high-variance regions into final DMR calls. Because seed-based expansion requires locally strong and directionally consistent signals, this strategy may limit the propagation of noisy windows during region construction. Together, these results indicate that seed-based integration complements variance-aware window detection by limiting the propagation of noisy window-level signals into region-level DMR calls.

### glmmDMR outperforms existing DMR detection methods across multiple benchmarking criteria

To evaluate how well glmmDMR performs compared to established DMR detection tools, we compared glmmDMR (SS, MS, and HB configurations) to DSS [13, 14], methylKit [11], DMRfinder [15], MACAU2 [16], metilene [24], and Fisher’s exact test [8, 12] using the same simulated datasets described above. Metilene detected relatively few DMRs under the simulation settings used here and therefore contributed minimally to subsequent comparative analyses (**Fig. 4a**). Across multiple benchmarking criteria, including detection accuracy, region structural fidelity, and robustness across methylation effect sizes, glmmDMR consistently outperformed all existing methods tested here.

**Fig. 4.**
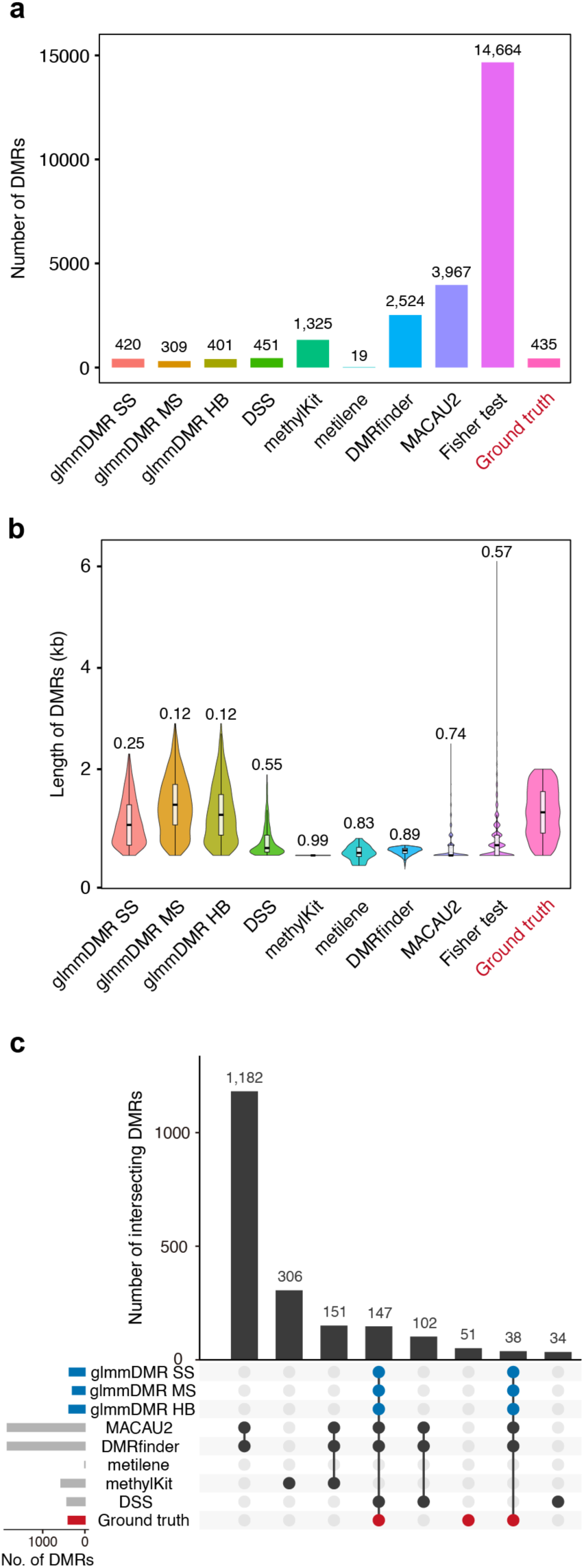
Benchmark comparison of glmmDMR and existing DMR detection methods using simulated methylome datasets. (**a**) Number of DMRs detected using different glmmDMR configurations (SS, single-seed; MS, multiple-seed; HB, hybrid) and existing DMR detection methods (DSS, methylKit, DMRfinder, MACAU2, and Fisher’s exact test). The number of ground-truth DMRs is shown in salmon (*n* = 435). (**b**) Distribution of DMR length (kb) for each method and for true simulated DMRs. Violin plots show the full distribution; boxplots show the median (center line) and IQR (box), with whiskers extending to 1.5× IQR. *D* values shown above each violin plot indicate the KS distance between the indicated distribution and the true simulated DMR length distribution. (**c**) Overlap analysis of DMRs detected by glmmDMR configurations and existing methods. The UpSet plot shows the number of DMRs in each intersection among methods. Only the eight largest intersections are shown for clarity.Bars indicate intersection sizes, and connected circles indicate the methods that contribute to each intersection. The red bar and red circles indicate the proportion of ground-truth DMRs within each intersection set. The blue bars indicate the number of DMRs detected by each of the three glmmDMR configurations.

Detection accuracy was consistently higher for glmmDMR across all configurations. PR analysis demonstrated that all three glmmDMR strategies maintain high precision across a broad range of recall values, whereas existing methods exhibited a rapid loss of precision with higher recall values (**Fig. S6a**). Fisher’s exact test achieved the high recall only at the expense of markedly reduced precision, indicating extensive FP detection. DMRfinder, MACAU2, and methylKit showed intermediate performance but failed to sustain precision at higher recall levels. Analysis of F1 scores confirmed that all glmmDMR configurations achieve a more favorable balance between sensitivity and specificity than the other evaluated methods (**Fig. S6c**).

The number of DMRs detected by glmmDMR was also more consistent with the ground-truth structure: glmmDMR produced region counts that closely matched the ground-truth number, whereas Fisher’s exact test markedly overestimated DMR counts, while the other methods also showed lower sensitivity by detecting more DMRs than expected (**Fig. 4a**). glmmDMR also more accurately reconstructed the structural properties of the true DMRs. The length distributions of the regions generated by glmmDMR closely matched those of the ground-truth DMRs, as evidenced by their relatively small KS distances from the ground-truth DMRs. By contrast, conventional approaches frequently generated fragmented or structurally inconsistent regions (**Fig. 4b**). Overlap analysis revealed that the largest intersections with the ground-truth DMRs involved the three glmmDMR strategies. By contrast, the overlaps between ground-truth DMRs and those detected by conventional methods were smaller (**Fig. 4c**). When plotting precision as a function of methylation effect size, we determined that all glmmDMR configurations maintain relatively high precision across a broad range of Δmethylation values, including at intermediate effect sizes where several existing methods showed marked declines in performance (**Fig. S6b**).

Consistent with our window-level and region-level findings, FP regions detected by all methods showed higher methylation variance among replicates than TP regions, whereas separation between TP and FP regions based on effect size was moderate and inconsistent across methods (**Fig. S7**). Together with the corresponding analyses for glmmDMR (**Fig. S5**), these results suggest that elevated methylation variance among replicates is a general feature of FP DMR calls across multiple analytical frameworks. The improved precision achieved by glmmDMR was accompanied by fewer high-variance regions among the detected DMRs.

glmmDMR required longer wall-clock time than simpler methods, reflecting the computational cost of replicate-aware mixed-model fitting and seed-based region integration (**Table 1**). However, its memory usage was moderate and comparable to that of existing tools. Given the gains in detection accuracy and structural fidelity demonstrated above, this computational trade-off represents a reasonable cost for improved DMR detection performance.

**Table 1.**
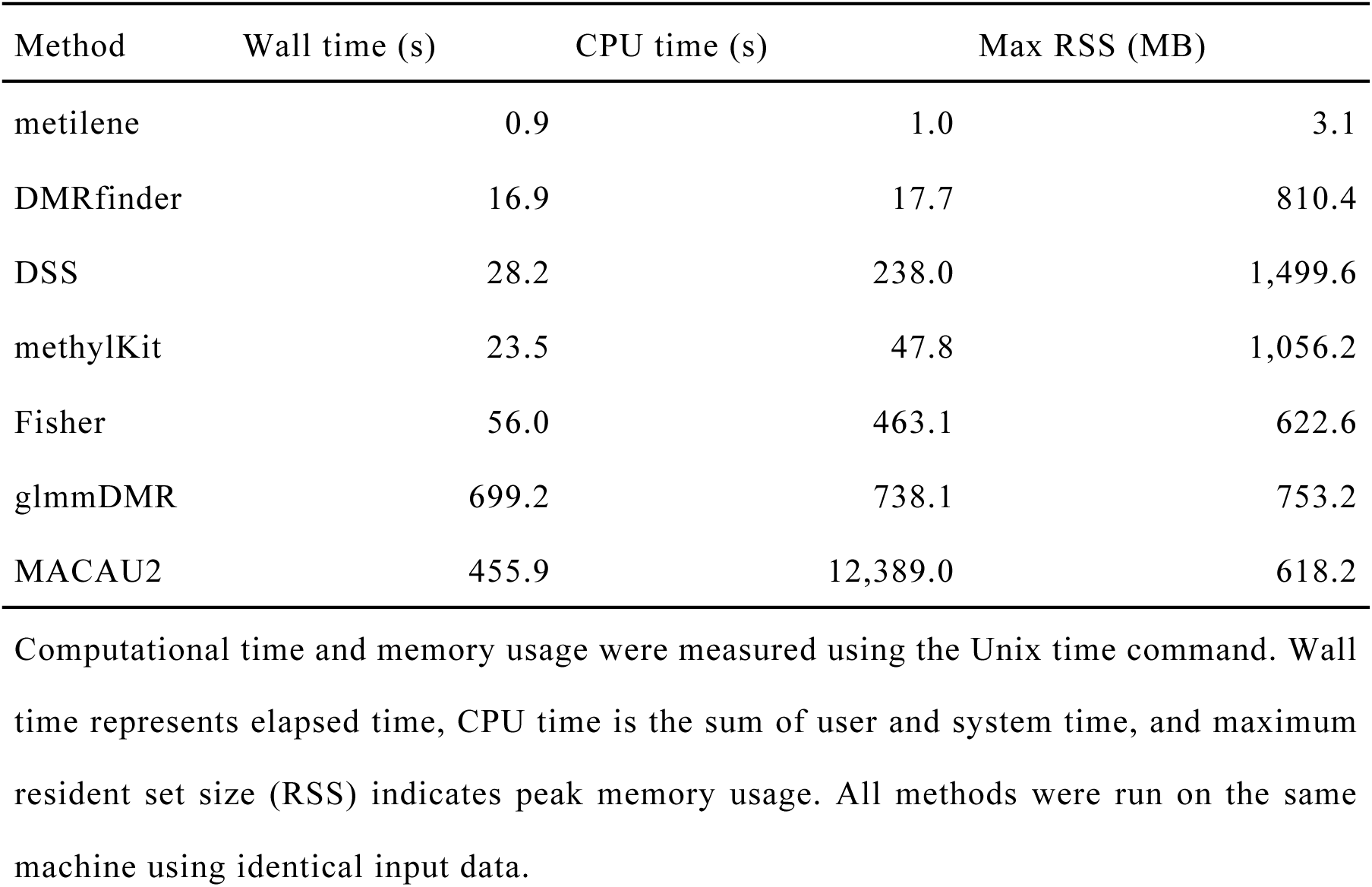
Computational performance of DMR detection methods.

### glmmDMR identifies biologically relevant DMRs in the *ddm1* mutant methylome

To evaluate glmmDMR under real biological conditions, we analyzed WGBS data from the wild-type Arabidopsis (*Arabidopsis thaliana*) accession Col-0 and the well-characterized *ddm1-2* mutant [25], which lacks function of the chromatin remodeler DECREASE IN DNA METHYLATION 1 (DDM1) and shows widespread DNA hypomethylation, predominantly at TEs [26, 27]. For this analysis, we compared glmmDMR (SS, MS, and HB configurations) to DSS and DMRfinder.

The evaluated methods differed substantially in the numbers and characteristics of the detected DMRs. glmmDMR identified 6,609, 5,011, and 7,795 hypomethylated DMRs under the SS, MS, and HB strategies, respectively, compared to 12,142 identified by DSS and 34,323 by DMRfinder (**Fig. 5a**). Hypomethylated regions (Hypo-DMRs) predominated across all methods, consistent with the global hypomethylation phenotype of *ddm1* mutants. glmmDMR produced longer regions than DMRfinder across all seed strategies. DMRfinder generated shorter, more fragmented regions (**Fig. 5b**), consistent with the differences in region-reconstruction behavior observed in the simulation analyses. The DMRs detected by glmmDMR also showed larger and more consistent differences in absolute methylation between groups than those identified by DSS or DMRfinder, which included a higher proportion of regions with smaller effect sizes (**Fig. 5c**). These patterns are consistent with the detection behavior observed in the simulation analyses.

**Fig. 5.**
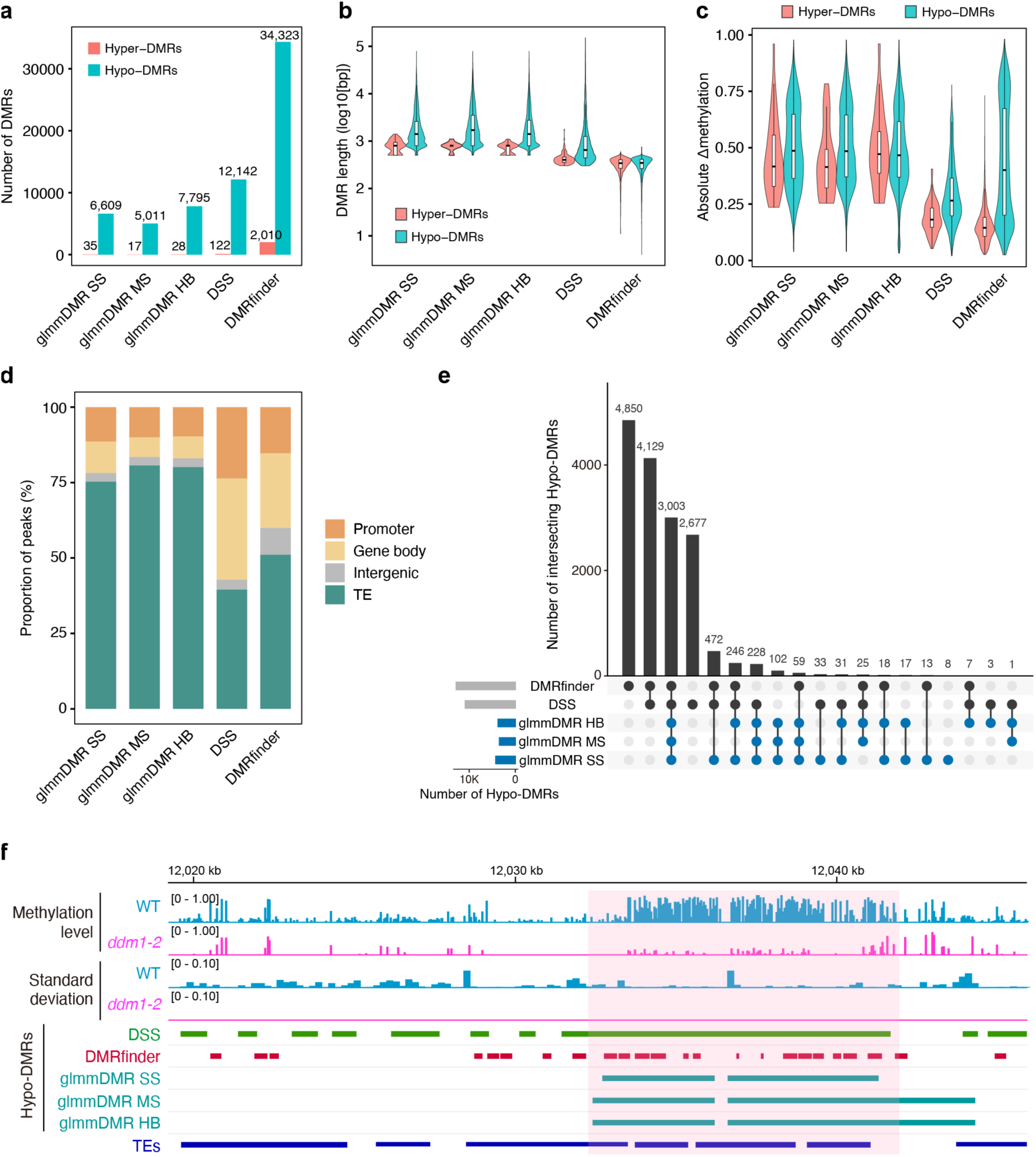
Comparison of DMR detection methods applied to the Arabidopsis *ddm1* methylome. (**a**) Number of hypermethylated DMRs (Hyper-DMRs, red) and hypomethylated DMRs (Hypo-DMRs, blue) detected by each method in the CG methylation context of the *ddm1* mutant. glmmDMR SS, MS, and HB represent the single-seed, multiple-seed, and hybrid strategies, respectively. (**b**) Distribution of DMR length (log10[bp]) for Hyper-DMRs and Hypo-DMRs detected by each method. Violin plots show the full distribution; boxplots show the median and IQR, with whiskers extending to 1.5× IQR. (**c**) Distribution of absolute methylation differences (|Δmethylation|) within detected DMRs for each method, stratified by DMR direction. Violin plots show the full distribution; boxplots show the median and IQR, with whiskers extending to 1.5× IQR. (**d**) Annotation of detected hypo-DMRs by genomic feature. Stacked bars show the proportions of DMRs that overlap with transposable elements (TEs), gene bodies, promoters, or intergenic regions for each method. (**e**) Overlap analysis of Hypo-DMRs detected in the CG context among glmmDMR configurations (SS, MS, HB), DSS, and DMRfinder. The UpSet plot shows the number of shared Hypo-DMRs in each intersection. The blue bars show the number of Hypo-DMRs detected by each of the three glmmDMR configurations. (**f**) Representative genomic locus showing CG methylation levels (top), the standard deviation of methylation levels between two biological replicates calculated for each genomic window (middle), and Hypo-DMR calls (bottom) for WT (blue) and *ddm1* (magenta) samples. Shaded regions indicate representative Hypo-DMRs detected by glmmDMR. TEs are shown as dark blue rectangles at the bottom.

Genomic annotation revealed that the Hypo-DMRs detected by glmmDMR are strongly enriched at TEs across all seed strategies, with substantially higher TE proportions than those observed for DSS or DMRfinder (**Fig. 5d**). This enrichment is consistent with the established role of DDM1 in maintaining transposon methylation and silencing [28, 29]. DSS and DMRfinder showed substantially lower enrichment of DMRs at TEs and correspondingly higher representation of non-TE features, suggesting broader, less specific detection. The DMRs identified by glmmDMR were highly reproducible across all three seed strategies, with the largest overlap consisting of 4,850 Hypo-DMRs shared by all three glmmDMR seed strategies (SS, MS, and HB only) (**Fig. 5e**). By contrast, DSS and DMRfinder each identified numerous regions that were not detected by glmmDMR, reflecting substantial differences in the regions prioritized by each method.

Metaplot analysis of methylation levels across DMR bodies revealed consistent differences in signal structure among the methods across all three methylation contexts (**Fig. S8**). Regions detected by DMRfinder showed sharp methylation transitions at the region boundaries, consistent with detection strategies based on local significance thresholds. By contrast, regions detected by glmmDMR exhibited more gradual methylation changes, with a peak at the center of the region body. This difference in signal shape is consistent with the seed-based expansion strategy of glmmDMR, which favors spatially consistent methylation domains over a series of locally significant windows. Inspection of representative genomic loci supported these observations (**Fig. 5f**). Regions detected by all methods showed clearly lower methylation levels in *ddm1-2* relative to the wild type. By contrast, DSS and DMRfinder each identified many additional regions not detected by glmmDMR. Although ground-truth DMRs are unavailable in real biological datasets and the true status of these regions therefore cannot be determined directly, their statistical characteristics are consistent with those of false-positive detections in the simulation analyses.

We observed broadly similar trends in the CHG and CHH methylation contexts, with glmmDMR producing more consistent and TE-enriched DMRs than DSS or DMRfinder. However, these contexts exhibited greater overall variability and smaller effect sizes (**Fig. S9**). Together, these results demonstrate that glmmDMR identifies biologically relevant and reproducible DMRs in real methylome datasets. The detected regions were highly enriched at TEs, showed high reproducibility across seed strategies, and exhibited methylation profiles consistent with the expected biological consequences of DDM1 loss.

### glmmDMR reveals TE-associated methylation changes in the *Osdml3a-1* mutant

Having examined the performance of glmmDMR on the well-characterized *ddm1-2* methylome, we asked whether it could detect subtler methylation changes in a mutant that lacked a clear developmental phenotype. *OsDML3a* encodes a putative DEMETER-like DNA glycosylase and its expression is upregulated during endosperm development (**Fig. S10a**). To investigate the role of OsDML3a in endosperm methylation, we generated the loss-of-function mutant *Osdml3a-1* in the Nipponbare background via CRISPR/Cas9-mediated gene editing (see Methods, **Fig. S10b**). Under standard growth conditions, *Osdml3a-1* plants showed no obvious developmental abnormalities, including in plant height, flowering time, panicle morphology, and endosperm development.

glmmDMR identified numerous DMRs in *Osdml3a-1* endosperm relative to wild-type endosperm across all three methylation contexts. In contrast to the predominantly hypomethylated profile observed in *ddm1-2*, hypermethylated DMRs (Hyper-DMRs) were predominant in *Osdml3a-1* (for the CG context: SS, 6,021; MS, 3,409; HB, 5,815 Hyper-DMRs versus SS, 2,580; MS, 1,640; HB, 2,530 Hypo-DMRs), consistent with loss of an active DNA demethylation function (**Fig. 6a**). We observed this pattern across the CG, CHG, and CHH contexts with all three seed strategies. The MS strategy detected fewer DMRs than the SS and HB strategies, consistent with its more stringent requirement for clustered evidence during seed selection, as noted during the simulation analyses. The three seed strategies produced DMRs with broadly similar length distributions and methylation effect size (**Fig. S11a,b**), indicating that differences among the strategies primarily affected the number of detected regions rather than their overall characteristics. Notably, most detected DMRs exhibited relatively modest methylation differences between groups, suggesting that the *Osdml3a-1* methylome is characterized by subtle but reproducible methylation changes rather than large-scale disruption of the methylome.

**Fig. 6.**
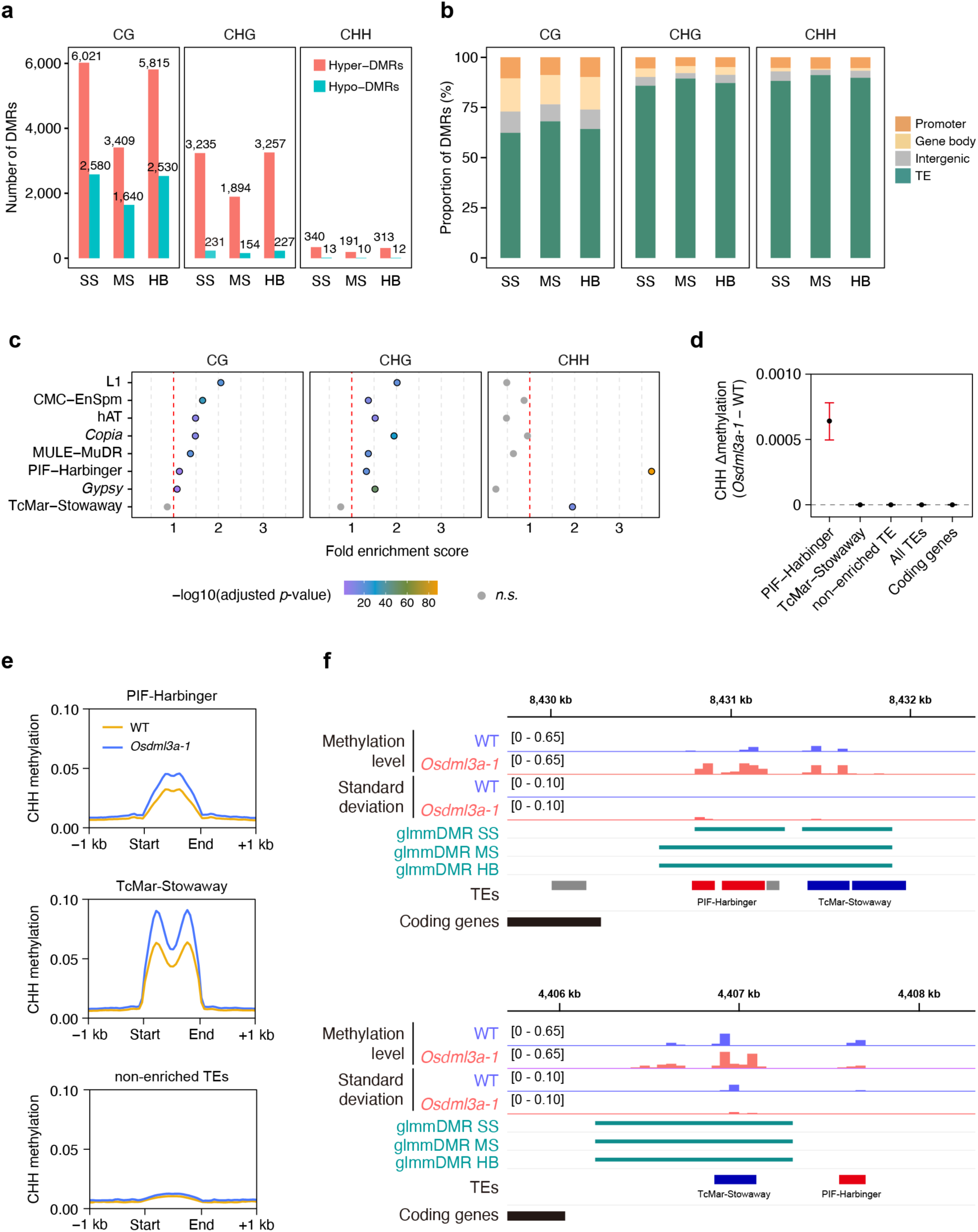
glmmDMR identifies subtle TE-associated methylation changes in the endosperm of the *Osdml3a-1* mutant. (**a**) Numbers of Hyper-DMRs (salmon) and Hypo-DMRs (blue) detected by glmmDMR in the CG, CHG, and CHH methylation contexts in the *Osdml3a-1* endosperm relative to those of the wild-type Nipponbare. SS, MS, and HB indicate the single-seed, multiple-seed, and hybrid strategies, respectively. (**b**) Genomic annotations of Hyper-DMRs detected in each methylation context. Stacked bars show the proportions of Hyper-DMRs that overlap with TEs, gene bodies, promoters, or intergenic regions for each seed strategy. (**c**) Statistical enrichment of TE families among Hyper-DMRs detected by glmmDMR using the HB strategy. The bubble plots show fold enrichment, with colors indicating −log10(adjusted *p*-value) calculated by the hypergeometric test. The red dashed vertical lines indicate fold enrichment > 1. Results from individual seed strategies are shown in Supplementary **Fig. S11c**. (**d**) Family-wide methylation differences across representative TE families. Points indicate median Δmethylation (*Osdml3a-1* − wild type) calculated using all annotated copies of each TE family; error bars indicate 95% bootstrap confidence intervals. Gene regions, all TEs combined, and non-enriched TE families are included as controls. Positive values indicate higher methylation in *Osdml3a-1* relative to Nipponbare. (e) Metaplots showing CHH methylation levels across the bodies of enriched (PIF-Harbinger and TcMar-Stowaway) and non-enriched TE families in wild-type (orange) and *Osdml3a-1* (blue) endosperm. The *x*-axis represents the relative distance between the TE start and end coordinates, scaled to a normalized body length, with 1-kb flanking regions on each side. (f) Representative genomic loci, showing CHH methylation levels (top), replicate-level standard deviation of CHH methylation levels (middle), and Hyper-DMR calls detected using the SS, MS, and HB strategies (bottom) in wild-type (WT) and *Osdml3a-1* endosperm. The upper locus is associated with a PIF-Harbinger element (red) and the lower locus with a TcMar-Stowaway element (blue). TE coordinates are shown below each locus, and coding genes are indicated with black boxes.

Genomic annotation of the Hyper-DMRs revealed strong enrichment at TEs across all methylation contexts and seed strategies (**Fig. 6b**). TE enrichment analysis identified specific TE families as being preferentially associated with these Hyper-DMRs (**Fig. 6c**). Multiple TE families, including L1, CMC-EnSpm, hAT, *Copia*, and MULE-MuDR, showed significant enrichment in the CG and CHG contexts. By contrast, enrichment in the CHH context was restricted to a small number of TE families, most notably PIF-Harbinger and TcMar-Stowaway. Similar enrichment patterns were observed across all three seed-based strategies (**Fig. S11c**).

To assess whether these enrichment patterns reflected methylation changes shared across entire TE families, rather than being restricted to DMRs, we quantified family-wide methylation differences between groups using all annotated copies of each TE family (**Fig. 6d; Fig. S11d**). Methylation in the CG and CHG contexts showed broadly positive but relatively small shifts in enrichment across multiple TE families. By contrast, high family-wide CHH methylation was much more restricted and was most evident in PIF-Harbinger elements (**Fig. 6d**). Although TcMar-Stowaway elements were significantly enriched among the CHH Hyper-DMRs (**Fig. 6c**), this family-wide methylation shift was comparatively weak (**Fig. 6d**), suggesting that methylation changes in this family may be confined to a subset of elements. Consistent with these observations, CHH metaplots revealed elevated methylation across PIF-Harbinger elements and localized higher methylation within TcMar-Stowaway elements in *Osdml3a-1* (**Fig. 6e; Fig. S11e,f**). An examination of representative genomic loci supported these observations. At both PIF-Harbinger- and TcMar-Stowaway-associated regions, CHH Hyper-DMRs were consistently detected by all three seed-based strategies and coincided with localized higher CHH methylation levels in *Osdml3a-1* (**Fig. 6f**).

Together, these results demonstrate that loss of OsDML3a function is associated with reproducible hypermethylation of specific TE families in the rice endosperm. Whereas higher CG and CHG methylation signals were broadly distributed across multiple TE families, changes in CHH methylation were concentrated to a limited subset of families, particularly PIF-Harbinger elements. These findings suggest that OsDML3a-dependent demethylation is not uniformly distributed across the endosperm methylome but preferentially affects specific classes of TEs.

## Discussion

This study was motivated by a central challenge in DMR detection methodology: why do methods with apparently comparable statistical power differ substantially in their FP behavior? Our analyses suggest that, under conditions of realistic methylome variability, spurious DMR detections are more strongly associated with high variability in replicate-level methylation than with small effect sizes alone. This finding has direct implications for the design and evaluation of DMR detection frameworks, as it indicates that sensitivity to effect size is insufficient for the reliable identification of biologically meaningful methylation changes between groups.

Our finding that methylation variability among biological replicates is a major contributor to FP detection reframes an important aspect of DMR analysis. Conventional benchmarking has often emphasized the ability to detect small methylation differences, which remains an important criterion [17, 18]. However, our results suggest that high-variance loci are an additional, often underappreciated source of spurious DMR calls. DNA methylation data are characterized by substantial overdispersion and biological variability among replicates, features that simple statistical models often fail to capture adequately [9, 10]. The GLMM framework implemented in glmmDMR addresses this issue by incorporating biological replicates as random effects [16, 19], enabling the direct modeling of replicate-level variability during statistical testing. This formulation improves discrimination between consistent methylation differences across biological groups and unstable high-variance signals among biological replicates, at least under the simulated conditions tested here. These findings also suggest that a fixed methylation-difference threshold may not always provide the most informative criterion for DMR detection. Because replicate-level methylation variance was more strongly associated with false-positive detections in simulations, and regions exhibiting high replicate-level variability were preferentially detected by conventional methods in both the Arabidopsis *ddm1-2* and rice *Osdml3a-1* methylomes, variance-aware statistical modeling may provide a useful alternative to approaches that rely heavily on predefined Δmethylation cutoffs.

A second contribution of glmmDMR lies in its seed-based region-construction strategy, which addresses a separate problem from variance-aware window-level testing. Biologically meaningful methylation changes often span contiguous genomic regions but may contain locally weak or non-significant windows due to coverage heterogeneity, variation in cytosine-site density across genomic regions, or stochastic sampling effects inherent to WGBS and EM-seq data. When DMRs are defined by directly merging individual significant windows, such interruptions cause true DMRs to be fragmented into multiple shorter regions, and peripheral noisy windows may be incorporated into the final call set. By initiating region construction from local high-confidence seeds and allowing directional extension based on regional signal consistency rather than requiring every constituent window to individually exceed a significance threshold, glmmDMR reconstructs DMRs that more closely reflect underlying biological methylation patterns while maintaining statistical support throughout the region. This signal-interruption tolerance distinguishes seed-based integration from simple window-merging approaches and may be particularly important for genomic features such as TEs and gene bodies, in which the underlying biological unit of methylation regulation often spans the full length of the element rather than isolated subregions. The differences in signal shape observed in the metaplot analyses, namely sharp boundary-associated transitions in DMRfinder, weaker boundary effects in DSS, and smoother body-centered profiles in glmmDMR, are consistent with these differences in region-construction logic and suggest that the DMRs detected by glmmDMR may better capture the gradual nature of underlying methylation transitions. Together, these results indicate that robust DMR detection requires both accurate variance modeling and biologically informed region reconstruction.

Although glmmDMR maintained relatively high precision across intermediate methylation effect sizes, its performance reflects an inherent trade-off between sensitivity and specificity. By explicitly modeling replicate-level variability and requiring local consistency during region expansion, glmmDMR preferentially retains methylation differences between groups that are both statistically supported and reproducible across replicates. This conservative behavior may be advantageous in applications where precision is important, such as the prioritization of DMRs for downstream experimental validation. At the same time, users who wish to prioritize sensitivity may adjust the significance and expansion parameters according to their study design.

The application of glmmDMR to real methylome data supported the utility of this approach. In the well-characterized Arabidopsis *ddm1* mutant, glmmDMR identified DMRs that were consistent across seed strategies, enriched at TEs [28, 29], and characterized by more consistent methylation profiles than those detected by the comparison methods. Regions uniquely detected by DSS or DMRfinder often showed low methylation levels and high variability in wild-type samples, resembling the high-variance FP patterns observed in simulations. Although a definitive ground truth is unavailable for real data, these patterns are consistent with the simulation findings and suggest that the statistical properties observed under controlled conditions may translate into biologically relevant differences in real methylome analyses.

Application of glmmDMR to the *Osdml3a-1* mutant illustrated the ability of glmmDMR to detect subtle, context-specific methylation changes. *OsDML3a* encodes a putative DEMETER-like (DML) DNA glycosylase orthologous to Arabidopsis enzymes involved in locus-specific active demethylation [30]. Whereas the DML protein REPRESSOR OF SILENCING 1 (OsROS1a) has been identified as a major demethylase in rice reproductive tissues [30–32], the *Osdml3a-1* mutant provides a tractable system because it lacks clear developmental defects. In this mutant, glmmDMR detected moderate but reproducible hypermethylation at specific TE families across multiple methylation contexts. Although enrichment analyses identified several TE families associated with Hyper-DMRs, family-wide methylation analyses revealed a more nuanced pattern. Higher CG and CHG methylation signals were broadly distributed across multiple TE families and were generally modest in magnitude, whereas changes in CHH methylation were concentrated in a smaller subset of TE families. In particular, PIF-Harbinger elements showed consistent family-wide rises in CHH methylation levels, and TcMar-Stowaway elements exhibited significant enrichment among Hyper-DMRs but comparatively weak family-wide methylation shifts. This distinction suggests that DMR enrichment and family-wide methylation remodeling do not necessarily coincide and may reflect different modes of epigenetic regulation. One possible explanation for the preferential methylation changes observed in PIF-Harbinger elements is that these loci are particularly sensitive to disruption of OsDML3a-mediated demethylation during endosperm development. Whether OsDML3a recognizes these loci through sequence-specific mechanisms or acts more broadly through chromatin occupancy, as proposed for ROS1 [33], remains to be determined. More generally, these results suggest that OsDML3a contributes to the maintenance of low methylation levels in a restricted subset of TE families rather than functioning as a global regulator of endosperm methylation. These findings illustrate how variance-aware DMR detection can reveal structured epigenetic alterations that are difficult to distinguish from noise using conventional approaches. The methylation changes observed in *Osdml3a-1* were considerably more subtle than those observed in *ddm1-2* but were reproducible across biological replicates and seed-based detection strategies. Independent validation will be important, given the absence of prior methylome characterization for this rice mutant. Nevertheless, these results demonstrate the utility of variance-aware DMR detection for the identification of biologically meaningful methylation changes in systems with modest effect sizes and minor developmental phenotypes.

Several limitations should be considered. First, replicate-aware mixed-model fitting is more computationally demanding than simpler approaches, although memory usage remained practical in our analyses. Second, our benchmarking focused primarily on bulk WGBS and EM-seq data. Although glmmDMR accepts ONT-derived methylation calls via Dorado output, which are converted to the same per-site count format as WGBS data, systematic benchmarking using long-read data across diverse genomic contexts is an important future direction. Third, extension to single-cell methylome data [34] would require a different modeling strategy because cell-to-cell heterogeneity cannot be treated in the same way as variation among biological replicates. Finally, although we validated glmmDMR using plant methylomes, further evaluation with mammalian methylomes will be important because methylation density, genomic distribution, and sources of variability differ substantially among organisms.

## Conclusions

glmmDMR provides a replicate-aware framework for DMR detection that combines GLMM-based variance modeling with seed-based region construction. A key finding of this study is that the false-positive rate of DMR detection is more strongly associated with elevated variance in replicate-level methylation than with small effect sizes, identifying variance among replicates as an underappreciated but important determinant of DMR detection reliability. glmmDMR improves precision by explicitly modeling replicate-level variability as a random effect within the GLMM framework, while its seed-based region-construction strategy reduces artificial fragmentation and improves the recovery of biologically interpretable DMRs across simulated and real methylome datasets. Application of glmmDMR to methylome data from *ddm1-2* in Arabidopsis and *Osdml3a-1* in rice demonstrated that it can reliably detect biologically meaningful changes in methylation, including subtle TE-associated methylation changes in the *Osdml3a-1* mutant. Together, these findings suggest that robust DMR detection requires both accurate variance modeling and biologically informed region reconstruction and that both replicate-level methylation variability and effect size should be considered when designing and evaluating DMR detection methods. More broadly, variance-aware statistical modeling combined with biologically informed region construction provides a practical framework for improving the reliability, interpretability, and biological relevance of DMR detection across diverse methylome studies.

## Methods

### Simulation of synthetic methylation data

To benchmark DMR detection methods under realistic conditions, synthetic CG methylation datasets were generated using a custom R script (simulate_sites.R; https://github.com/ktonosaki/glmmDMR). Simulations were restricted to the CG context because the objective was to benchmark statistical model performance rather than context-specific methylation biology. A 10-Mb pseudo-chromosome was simulated, with cytosine positions generated following a Poisson distribution (*λ* = 20,000 sites per Mb). Two groups were simulated (wild type [WT] and mutant [MT]), each with four biological replicates.

DMRs were embedded in 5% of the genome as contiguous blocks of 300–2,000 bp, with Δmethylation values ranging from 0.1 to 1.0 at intervals of 0.1. Each DMR was randomly designated as hypermethylated or hypomethylated in the MT samples. Within-DMR signal heterogeneity (heterogeneity among cytosine sites within each DMR) was introduced by restricting methylation shifts to a random subset of sites within each block, reflecting the incomplete methylation changes commonly observed in biological systems. Biological variability was modeled using a beta-binomial framework [9, 13] with the overdispersion parameter *ρ* = 0.3. Read coverage was sampled from a log-normal distribution (*μ* = 3.0, *σ* = 1.2 on the natural-log scale). A 30% missing rate was applied, and sites with fewer than five reads were excluded. Full simulation parameters are provided in Supplementary Table S1.

### GLMM for replicate-aware detection of differentially methylated windows

Before statistical modeling, the genome was tiled into 300-bp sliding windows with a step size of 200 bp. Windows that contained fewer than three cytosine sites after filtering were excluded from the analysis. To detect differentially methylated windows while explicitly accounting for biological variability across replicates, methylation data were modeled using GLMMs. Two dimensions of model specification were considered: data representation (site level vs. aggregated) and response distribution (binomial vs. beta), resulting in four model configurations.

In the binomial site–level model, methylated read counts *y_is_*_w_ out of total reads *n_is_*_w_ for site *s*, replicate *i*, and window *w* were modeled as:

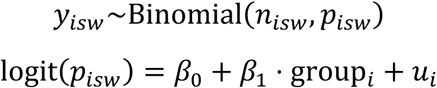

In the beta site–level model, methylation proportions *m_is_*_w_ were modeled as:

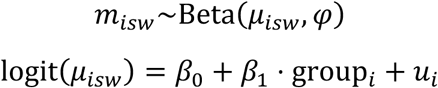

In the binomial-aggregated model, total methylated read counts *y_i_*_w_ and total coverage *n_i_*_w_ across sites within each window were used:

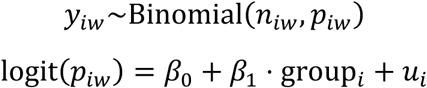

In the beta-aggregated model, window-level mean methylation proportions *m_i_*_w_

were modeled as:

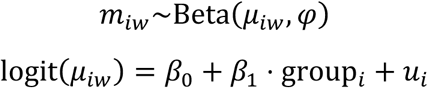

In all four configurations described above, *β*_0_ is the intercept, *β*_1_ is the fixed-effect coefficient for the group (WT vs. MT) on the logit scale, and *u_i_* ∼ *N*(0, *σ*^2^) represents replicate-specific random intercepts that capture biological variability among replicates. The variance parameter *σ*^2^ was estimated jointly with other model parameters during maximum likelihood optimization. In beta-regression models, the precision parameter *φ* was estimated during model fitting rather than fixed a priori, enabling the model to adaptively capture heterogeneity in methylation proportions across windows.

The statistical significance of the group effect (*β*_1_) was assessed with a likelihood ratio test comparing the full model to a reduced model without the group fixed effect. *β*_1_ represents the estimated group difference on the logit scale and is distinct from Δmethylation, which was computed independently as the difference in mean methylation proportions between groups on the response scale and used for descriptive and visualization purposes. *p*-values were adjusted for multiple testing using the Benjamini–Hochberg (BH) method [35]. All models were fitted using the glmmTMB package (v1.1.12) [19] in R (v4.4.2).

### Integration of differentially methylated windows into DMRs

To integrate window-level GLMM statistics into DMRs, a unified region-construction framework was implemented, consisting of three seed-based strategies used in glmmDMR and three conventional region-construction approaches used for benchmarking. All methods operated on identical window-level GLMM output and considered only windows that shared the same methylation direction.

### Seed-based region-construction strategies

For all seed-based strategies, window-level *p*-values within candidate regions were combined using the Stouffer method [20]. Each *p*-value (*p_i_*) was converted to a *z*-score (*z_i_* = Φ^−1^(1 − *p_i_*)). The regional test statistic was calculated as:

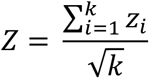

where *k* is the number of windows. The combined *p*-value was then computed as:

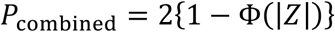

This approach weighs each window equally and yields a combined *p*-value that reflects both the number and consistency of significant windows within a region. Before region construction, all windows were assigned a methylation direction based on the sign of the estimated group effect (*β*_1_): windows with positive *β*_1_ were designated as hypermethylated and those with negative *β*_1_ as hypomethylated. Only windows that shared the same direction were considered for integration into a common region. The following three seed-based strategies constitute the core DMR-construction framework of glmmDMR and were benchmarked using identical input data and shared core parameters (Supplementary Table S2).

#### Single seed (SS)

Individual windows with *p* < 0.05 were designated as seeds. Starting from each seed, a region was expanded by iteratively incorporating adjacent windows with the same direction of methylation change (hyper- or hypomethylation). At each step, the Stouffer-combined *p*-value was recomputed for the growing region, and a candidate window was incorporated only if the updated combined *p*-value met the criteria listed in Supplementary Table S2. Expansion proceeded iteratively upstream and downstream of the starting seed until no qualifying windows remained.

#### Multiple seed (MS)

Seeds were defined at the cluster level. Neighboring same-direction windows within a 200-bp region were grouped into clusters, and a cluster was designated as a seed if its Stouffer-combined *p*-value fell below 0.05. Region expansion, including iterative re-computation of the Stouffer *p*-value, followed the same criteria as in the SS strategy.

#### Hybrid (HB)

The MS strategy was first applied across the genome to identify seed regions. The SS strategy was then applied independently to genomic regions not covered by any MS-derived seed. The resulting regions were combined into a single call set without additional merging.

### Conventional region-construction approaches used for benchmarking

For comparison, three conventional region-construction approaches were implemented using the same window-level GLMM output.

#### Simes aggregation

Window-level *p*-values within candidate regions were combined using the Simes test [23]. Candidate regions were defined by grouping windows with *p* < 0.05 within a 200-bp genomic region without seed-based expansion, and regions were retained at a combined *p*-value threshold of 0.0015.

#### Stouffer aggregation

Window-level *p*-values were combined using the Stouffer method [20] as described above. Candidate regions were defined by grouping neighboring windows with *p* < 0.05 within a 200-bp genomic region without seed-based expansion.

#### FDR-based merging

Windows with a BH-adjusted *p* < 0.05 were merged into contiguous regions within a 200-bp genomic region without *p*-value combination. Merged regions were retained if at least 50% of the constituent windows had BH-adjusted *p* < 0.05.

All region-construction strategies and benchmarking approaches were implemented using a custom R script (merge_window.R; https://github.com/ktonosaki/glmmDMR) and applied to identical window-level GLMM outputs.

### Benchmarking against existing DMR detection methods

For evaluation against established methods, glmmDMR was compared to DSS [13, 14], methylKit [11], DMRfinder [15], metilene [24], MACAU [16], and Fisher’s exact test using the same simulated methylation datasets. No minimum methylation-difference threshold was applied to any method to evaluate intrinsic statistical detection performance independently of user-defined effect-size cutoffs.

DSS (v2.52.0) was applied at the site level with local smoothing (smoothing.span = 500). DMRs were called with a *p*-value threshold of 0.05, a minimum of three CG sites per region, and a maximum merging gap of 200 bp.

methylKit (v1.3.33) was applied using the same sliding-window scheme as glmmDMR (300-bp windows, 200-bp step), and differential methylation was tested at *q* < 0.05. Adjacent significant windows were merged using a maximum merging gap of 200 bp.

DMRfinder (v0.3) was run with default parameters, except that no minimum methylation difference was applied (-d 0) and a minimum of three CG sites per region was required.

metilene (v0.2-9) was included as a representative segmentation-based DMR detection method and was run with a maximum gap of 200 bp, a minimum of three CG sites per DMR, and no minimum methylation difference threshold, followed by post-processing with metilene_output.pl.

MACAU2 (v2.0) was applied at the window level using the binomial mixed model with an identity-relatedness matrix, which was appropriate because the simulated samples lack population structure. Significant windows (*p* < 0.05) within a 200-bp region were merged into DMRs.

Fisher’s exact test was applied at the window level by aggregating read counts across replicates within each group. *p*-values were adjusted using the BH method, and significant windows (false discovery rate [FDR] < 0.05) within a 200-bp region were merged.

### Performance evaluation metrics

Detection performance was evaluated at both the window and DMR levels using a common set of benchmarking metrics. At the window level, each tested window was classified as a true positive (TP) if it overlapped with a ground-truth DMR by at least 1 bp and as a false positive (FP) otherwise. At the DMR level, a detected region was classified as a TP if it overlapped with a ground-truth DMR coordinate by at least 1 bp and as an FP otherwise.

Precision–recall (PR) curves and receiver operating characteristic (ROC) curves were generated by varying the statistical significance threshold applied to window-level *p*-values or DMR-level combined *p*-values, depending on the method being evaluated. Precision was defined as TP/(TP + FP) and recall as TP/(TP + FN), where FN denotes the number of false negatives. The area under the PR curve (AUPRC) and the area under the ROC curve (AUROC) were computed as summary statistics. F1 scores were calculated as the harmonic mean of precision and recall at each threshold, and the maximum F1 score across all thresholds was used as a single-value summary of overall detection performance. The structural accuracy of the detected DMRs was assessed by comparing the distributions of DMR length between each method and the ground-truth DMR set using the Kolmogorov–Smirnov (KS) statistic. Smaller KS distances indicate closer agreement with the true DMR length distribution. To characterize the features associated with true and spurious detections, methylation effect size (absolute Δmethylation), methylation variance across replicates, mean sequencing coverage, and coverage variance were compared between TP and FP windows or DMRs for each method. The statistical significance of differences was assessed using the Wilcoxon rank-sum test.

### Evaluation of computational performance

The computational performance of all methods was benchmarked on a single server equipped with two Intel Xeon Gold 6338 processors (64 physical cores, 128 threads in total; 2.00 GHz base clock) and 1 TiB RAM running Ubuntu 20.04.5 LTS (kernel 5.15.0-130). All methods were evaluated using the same simulated methylome dataset used for the benchmarking analyses.

Wall-clock time and peak memory usage (maximum resident set size, RSS) were measured using /usr/bin/time -v. The exact command-line options and thread counts used for each method are provided in Supplementary Table S3.

### Plant materials and generation of *Osdml3a-1*

The rice (*Oryza sativa* ssp. *japonica*) cultivar Nipponbare was used for all experiments. Plants were grown under short-day conditions (11-h light/13-h dark photoperiod with a 30°C/25°C day/night cycle). To generate *Osdml3a* mutants, a single guide RNA targeting exon 1 of *OsDML3a* (5′-AGCAATCCCGCTCCGTGTCT-3′) was cloned into pU6gRNA-oligo and inserted into pZH_OsU3gYSA_MMCas9 [36]. Rice transformation was performed using *Agrobacterium tumefaciens* strain EHA105. The *Osdml3a-1* mutant carries a 730-bp deletion spanning the upstream region and exon 1 of *OsDML3a*, together with a 13-bp insertion, and is predicted to disrupt OsDML3a function (**Fig. S10b**).

After backcrossing to wild-type Nipponbare for two generations to remove the transgene and minimize potential off-target effects, homozygous *Osdml3a-1* plants were identified by analysis of simple sequence length polymorphisms using primers flanking the deletion (F: 5′-TTGATGGATGTTGTGGATGG-3′, R: 5′-CTTTGTCGCCACACTCACTC-3′) and confirmed by agarose gel electrophoresis.

### RT-qPCR analysis

All tissues were collected from Nipponbare plants grown under short-day conditions. Samples included immature seeds at 4–7 days after pollination (DAP) and 10 DAP, leaves, roots, and spikelets collected from 4-week-old plants. Three independent biological replicates were analyzed. Total RNA was extracted using an RNeasy mini kit (QIAGEN) and reverse transcribed into cDNA using a PrimeScript kit (TaKaRa). Quantitative PCR (qPCR) was performed using a Thermal Cycler Dice Real Time System TP800 (TaKaRa) with gene-specific primer sets (Supplementary Table S4). Relative expression levels were normalized against *UBIQUITIN*, using primers that amplify both *LOC_Os03g13170* and *LOC_Os09g39500*.

### DNA extraction and EM-seq library preparation

Endosperm tissues were dissected from the seeds of BC2 homozygous *Osdml3a-1* and wild-type plants at 10 DAP (two biological replicates per genotype). Genomic DNA was extracted using a DNeasy kit (QIAGEN). EM-seq libraries were prepared using an NEBNext Enzymatic Methyl-seq Kit (New England Biolabs) according to the manufacturer’s instructions, then sequenced on a HiSeq X Ten instrument (Illumina) to generate 150-bp paired-end reads.

### Methylome data analysis

WGBS data from the rosette leaves of the wild type *Arabidopsis thaliana* accession Col-0 and the *ddm1-2* mutant [25] and EM-seq data from wild-type (Nipponbare) and *Osdml3a-1* endosperm were processed using the following pipeline. Raw reads from both datasets were quality-trimmed using fastp (v0.24.0) [37] with a minimum base quality of 30, sliding-window trimming (window size 4, mean quality 20), and a minimum read length of 36 bp. Trimmed reads were aligned to the *Arabidopsis thaliana* TAIR10 [38] or rice IRGSP-1.0 [39] reference genome using Bismark (v0.24.0) [40] with Bowtie2 [41]. Only uniquely mapped reads were retained, and PCR duplicates were removed using deduplicate_bismark. Methylation calls were extracted for the CG, CHG, and CHH contexts using bismark_methylation_extractor. Within each sample, sites with fewer than five reads were excluded from downstream analyses.

The outputs of Bismark methylation extractor were summarized into per-site methylated and unmethylated read counts using summarize_extractor.py. Per-site methylation significance was assessed using BinomTest.py based on the estimated non-conversion rate. Context-specific sliding-window matrices were generated using prepare_matrix.sh with a window size of 500 bp and a step size of 300 bp. A larger window size was used for real methylome datasets than for simulations (300-bp windows with a 200-bp step size) to improve signal robustness, given the higher biological and technical variability characteristic of real WGBS and EM-seq data. GLMM-based differential methylation testing was performed using run_glmmDMR.R with the beta site-level model and biological replicates modeled as random effects. DMRs were constructed from window-level statistics using merge_window.R, as described above. For comparison, DSS and DMRfinder were applied to the same datasets using the parameters described in the Benchmarking section.

Genomic features were defined using the TAIR10 GFF3 annotation file for Arabidopsis [42] and the MSU version 7.0 annotation for rice [39]. Promoters were defined as the 2-kb regions upstream of each annotated TSS. TE annotations were obtained from the transposon annotation in the TAIR10 GFF3 file for Arabidopsis and the GFF3 TE annotation available through the UCSC Table Browser of Plant cis-Map (http://ucsc.gao-lab.org) for rice. Intergenic regions were defined as genomic intervals that did not overlap with TEs, promoters, or gene bodies. DMRs were assigned to genomic features by hierarchical annotation using bedtools intersect (v2.30.0) [43]: DMRs that overlapped with TEs were assigned to the TE category first, followed by promoters, gene bodies, and intergenic regions. Per-sample methylation levels were summarized into 50-bp binned bigWig files using make_binned_methylation_bigwig.R for visualization, and metaplots were generated by scaling DMR bodies to a normalized length with 1 kb of flanking regions on each side. To evaluate whether methylation changes were shared across entire TE families, family-wide methylation differences were calculated using all annotated copies of each TE family rather than only those members overlapping with DMRs. For each TE family, methylation levels were averaged across all cytosine sites within annotated family members for each biological replicate, and the median methylation difference was calculated. Confidence intervals were estimated by bootstrap resampling of TE copies (1,000 iterations), and 95% confidence intervals were defined from the 2.5th and 97.5th percentiles of the bootstrap distribution.

### Use of large language models

ChatGPT (OpenAI) was used to improve the readability and language of the manuscript. The authors reviewed and edited all AI-generated text and take full responsibility for the manuscript and its content.

## Abbreviations

DMR: differentially methylated region
GLMM: generalized linear mixed model
TE: transposable element
WGBS: whole-genome bisulfite sequencing
EM-seq: enzymatic methyl sequencing
SS: single-seed
MS: multiple-seed
HB: hybrid
TP: true positive
FP: false positive
PR: precision–recall
ROC: receiver operating characteristic
KS: Kolmogorov–Smirnov
IQR: interquartile range
DAP: days after pollination
F1 score: harmonic mean of precision and recall.

## Declarations

### Ethics approval and consent to participate

Not applicable.

## Consent for publication

Not applicable.

## Availability of data and materials

The WGBS data from the wild type and *ddm1-2* mutant in *Arabidopsis thaliana* used in this study are publicly available at the NCBI Sequence Read Archive under BioProject PRJNA967123 (accession numbers SRR24421264, SRR24421266, SRR24421268, and SRR24421270). The EM-seq data from the wild type and the *Osdml3a* mutant generated in this study have been deposited at the DNA Data Bank of Japan under BioProject PRJDB42535 (accession numbers DRX1038158, DRX1038159, DRX1038160, and DRX1038161) and will be released upon publication. glmmDMR software and all analysis scripts used in this study are available at https://github.com/ktonosaki/glmmDMR under an MIT License.

## Author contributions

K.T. conceived and designed the study. K.T. developed the glmmDMR framework, designed and performed the simulation analyses, interpreted the results, and wrote the manuscript. Y.D. performed the experiments and analyzed the methylome datasets. M.U. generated experimental material and produced the EM-seq data. T.K. supervised the research. All authors read and approved the final manuscript.

## Acknowledgements

We thank Kaho Yamaguchi and Yuki Shimizu for technical assistance with plant cultivation and molecular experiments. We also thank Akemi Ono, Yuichi Fukuda, and Kazuki Asai for assistance with sample preparation. Computational analyses were performed using the computational resources of the Kihara Institute for Biological Research, Yokohama City University, Japan. We thank members of the Kihara laboratory for helpful discussions and comments on this work.

## Supplementary information

Additional file 1: Supplementary figures S1–S11. A single PDF file containing all supplementary figures referenced in the manuscript. Captions and legends for each supplementary figure are included within the file.

Additional file 2: Supplementary tables S1–S4. A compiled multi-tab Excel file containing all supplementary tables referenced in the manuscript.

