## Supplementary_figures_S1_S11 for "glmmDMR reveals replicate-level methylation variance as a major determinant of false-positive DMR detection"

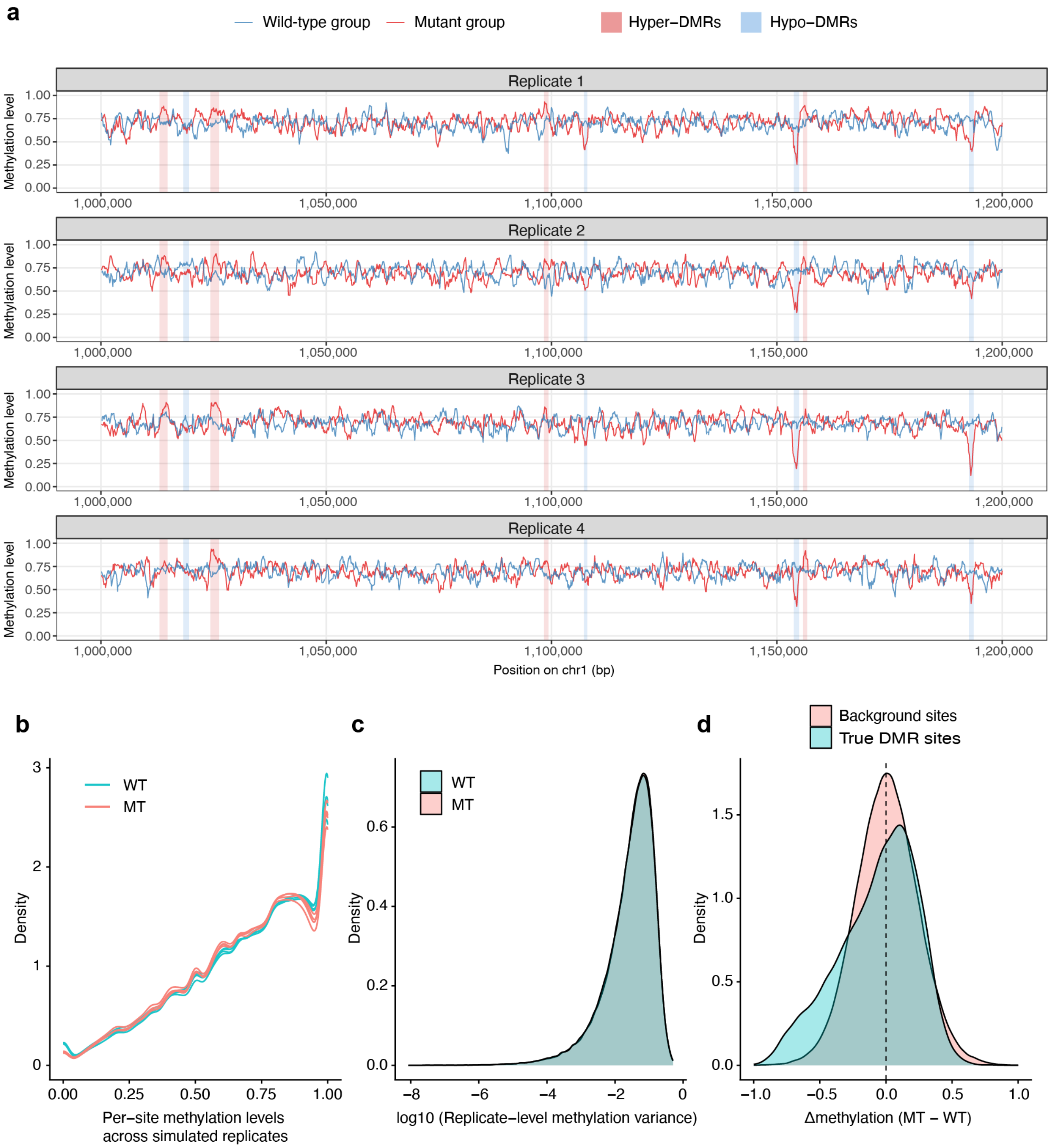


**Fig. S1 Representative simulated methylation profiles used for benchmarking analyses**
(**a**) Simulated methylation profiles across representative genomic regions for the wild-type and mutant groups. Each panel represents an independent simulated biological replicate. Blue and red lines indicate methylation levels for the wild-type and mutant groups, respectively. Shaded regions indicate simulated hypermethylated (red) or hypomethylated (blue) DMRs embedded within the background methylation landscape. The simulated datasets incorporate realistic local fluctuations in methylation levels, replicate-level variability, and heterogeneous effect sizes across genomic regions. (**b**) Distribution of per-site methylation levels across simulated biological replicates for the wild-type (WT) and mutant (MT) groups. (**c**) Distribution of per-site methylation variance among biological replicates for the WT and MT groups. (**d**) Distribution of mean methylation differences between groups (Δmethylation) for true DMR sites and background sites.

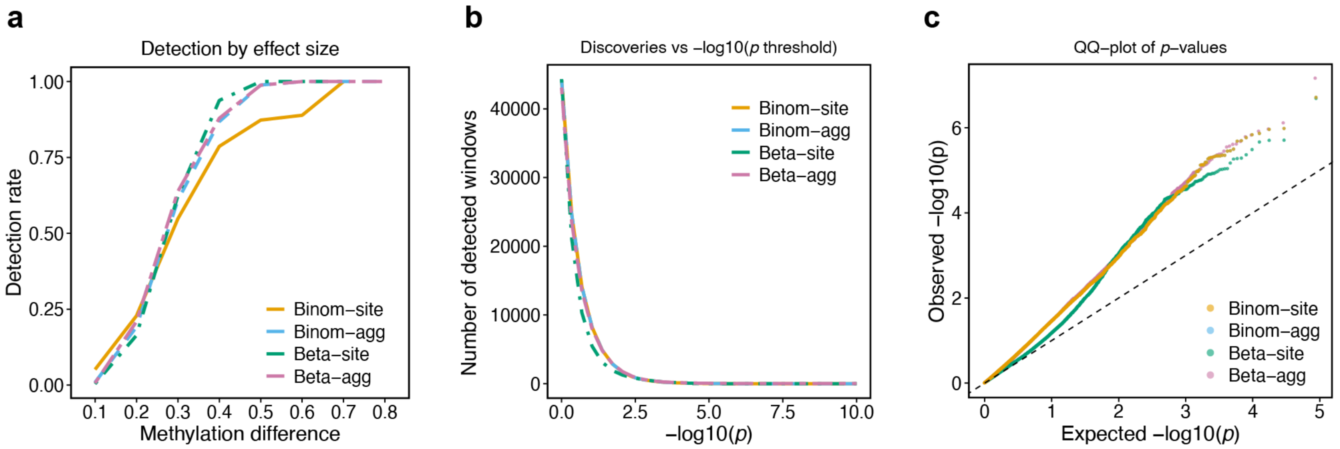


**Fig. S2 Comparison of GLMM model configurations using simulated methylome datasets**
(**a**) Detection rate (recall) as a function of methylation difference between groups (Δmethylation) for the four GLMM model configurations. (**b**) Number of detected windows across increasing statistical significance thresholds (–log10[*p*]). (**c**) Quantile–quantile (QQ) plot comparing expected and observed *p*-values for each GLMM model configuration. The dashed diagonal line indicates the expected distribution under the null hypothesis.

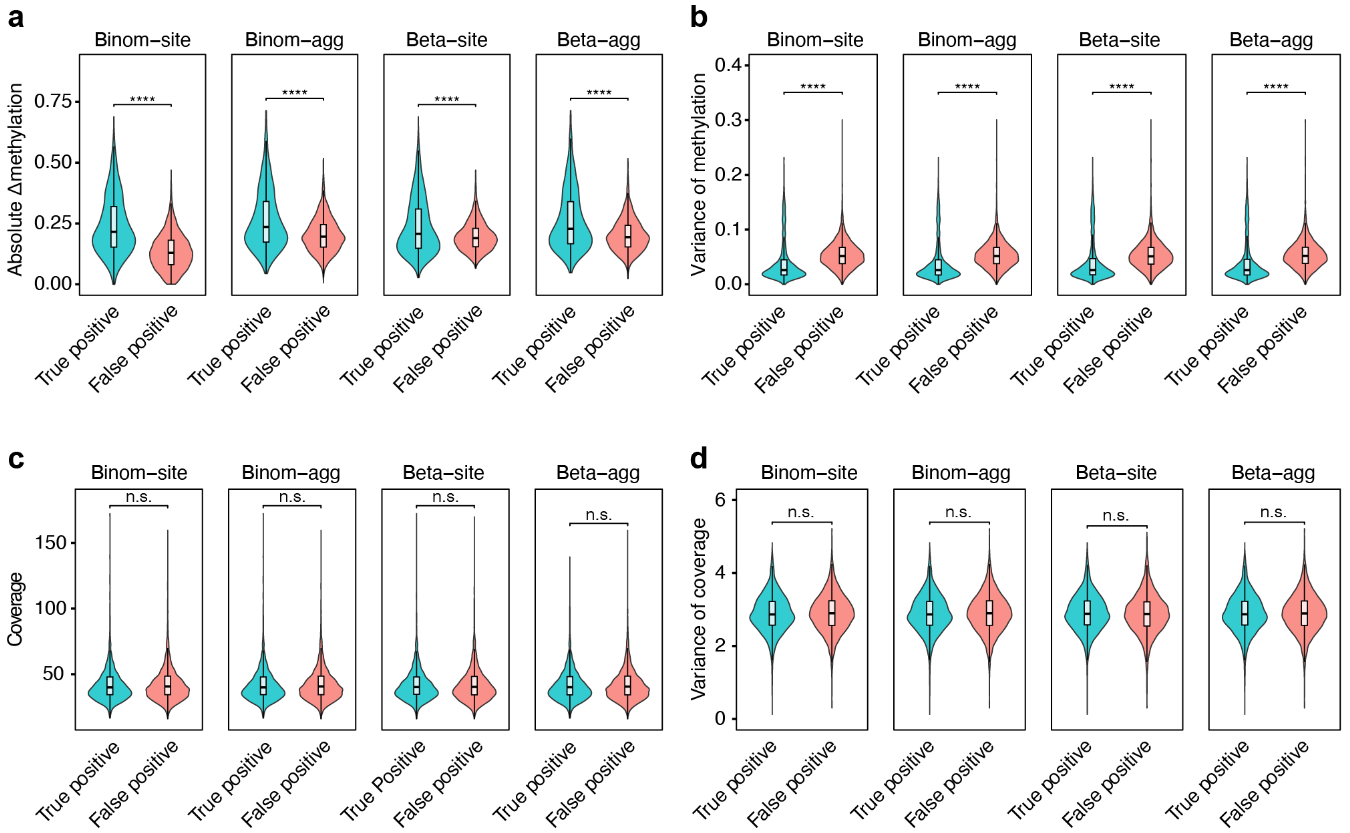


**Fig. S3 Characteristics of true-positive and false-positive windows across GLMM model configurations**

(**a, b**) Distribution of absolute methylation differences between groups. (**a**) and replicate-level methylation variance among replicates (**b**) for true-positive and false-positive windows across the four GLMM model configurations. (**c, d**) Distribution of sequencing coverage (**c**) and replicate-level variance in sequencing coverage (d) for true-positive and false-positive windows. Violin plots show the full distributions; boxplots show the median and interquartile range (IQR), with whiskers extending to 1.5× IQR. The significance of statistical differences between true-positive and false-positive groups was determined using the Wilcoxon rank-sum test (****$p < 0.0001$; ***$p < 0.001$; **$p < 0.01$; *$p < 0.05$; *n.s.*, not significant).

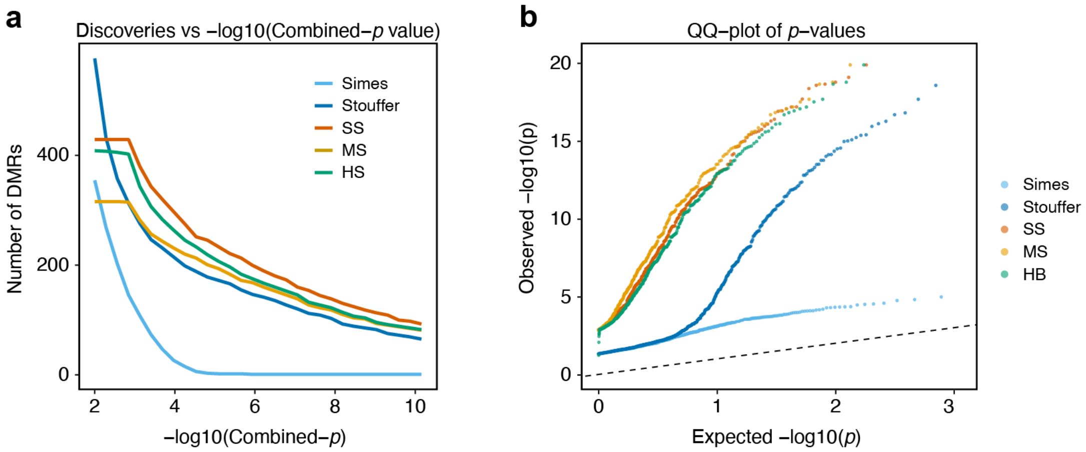


**Fig. S4 Comparison of seed-based DMR construction strategies**

(**a**) Number of detected DMRs across increasing combined statistical significance thresholds (–log10[combined *p*-value]) for seed-based and conventional region-construction methods. (**b**) QQ plot of expected and observed combined *p*-values for each DMR-construction strategy.

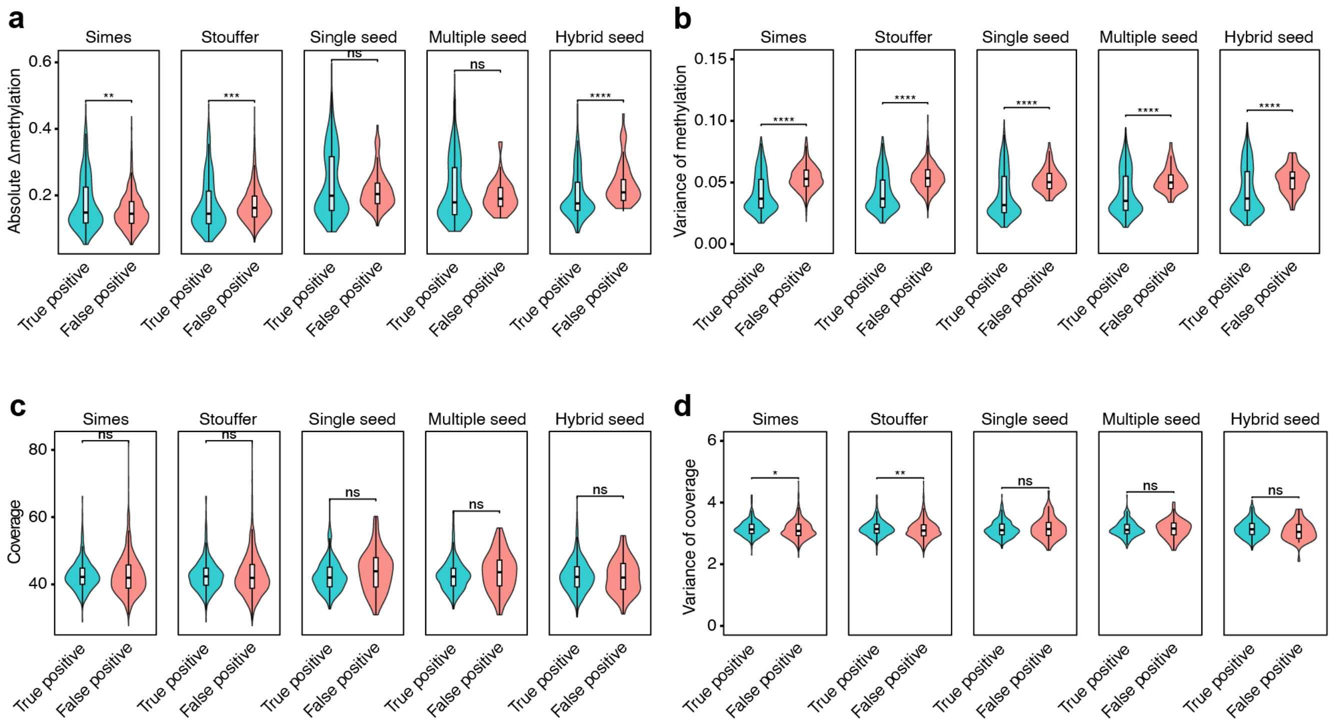


**Fig. S5 Characteristics of true-positive and false-positive DMRs across region-construction strategies**

(**a, b**) Distribution of absolute methylation differences between groups (**a**) and methylation variance among replicates (**b**) for true-positive and false-positive DMRs across different region-construction strategies. (**c, d**) Distribution of mean sequencing coverage (**c**) and coverage variance (**d**) for true-positive and false-positive DMRs. Violin plots show the full distributions; boxplots show the median and IQR, with whiskers extending to 1.5 × IQR. The significance of statistical differences between the true-positive and false-positive groups was determined using the Wilcoxon rank-sum test (****$p < 0.0001$; ***$p < 0.001$; **$p < 0.01$; *$p < 0.05$; *n.s.*, not significant).

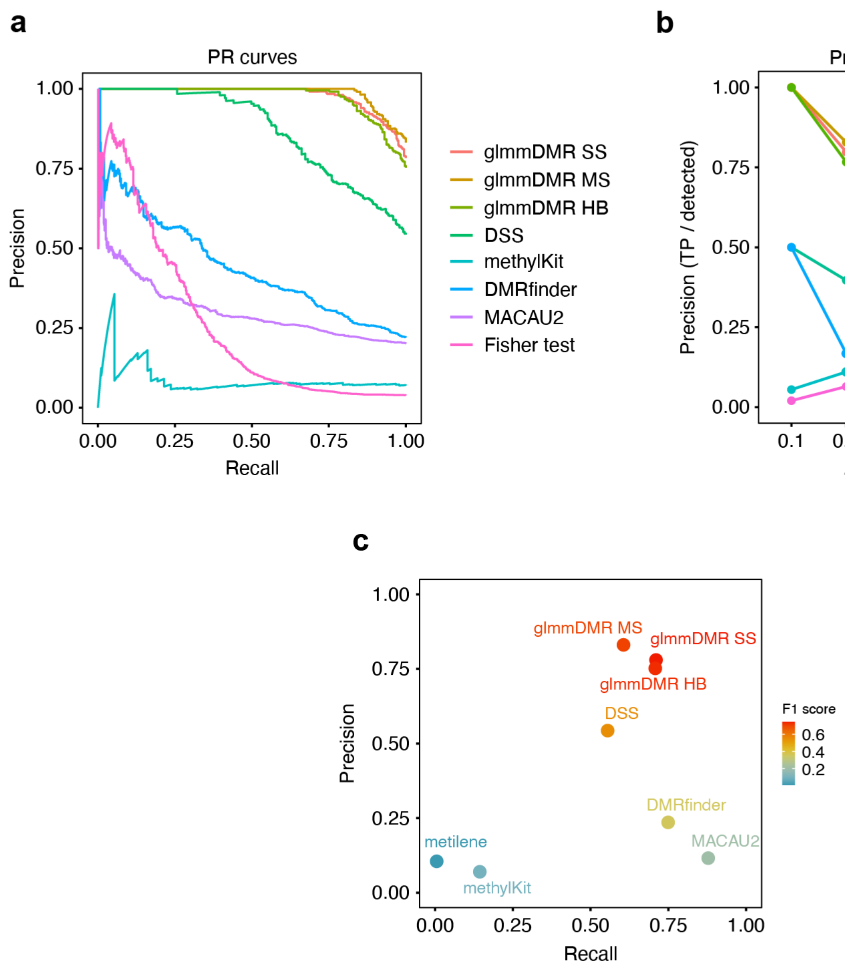


**Fig. S6 Benchmark comparison of glmmDMR with existing DMR detection methods**

(**a**) Precision–recall (PR) curves comparing the three glmmDMR configurations and existing DMR detection methods using simulated methylome datasets. glmmDMR SS, MS, and HB indicate single-seed, multiple-seed, and hybrid region-construction strategies, respectively. (**b**) Precision as a function of observed methylation difference. Precision was calculated separately for detected DMRs grouped by their observed absolute Δmethylation. Methods with no detected DMRs in a given bin are not shown for that bin. (**c**) Scatterplot showing the extent of correlation between recall and precision for each DMR detection method. Point color indicates the F1 score.

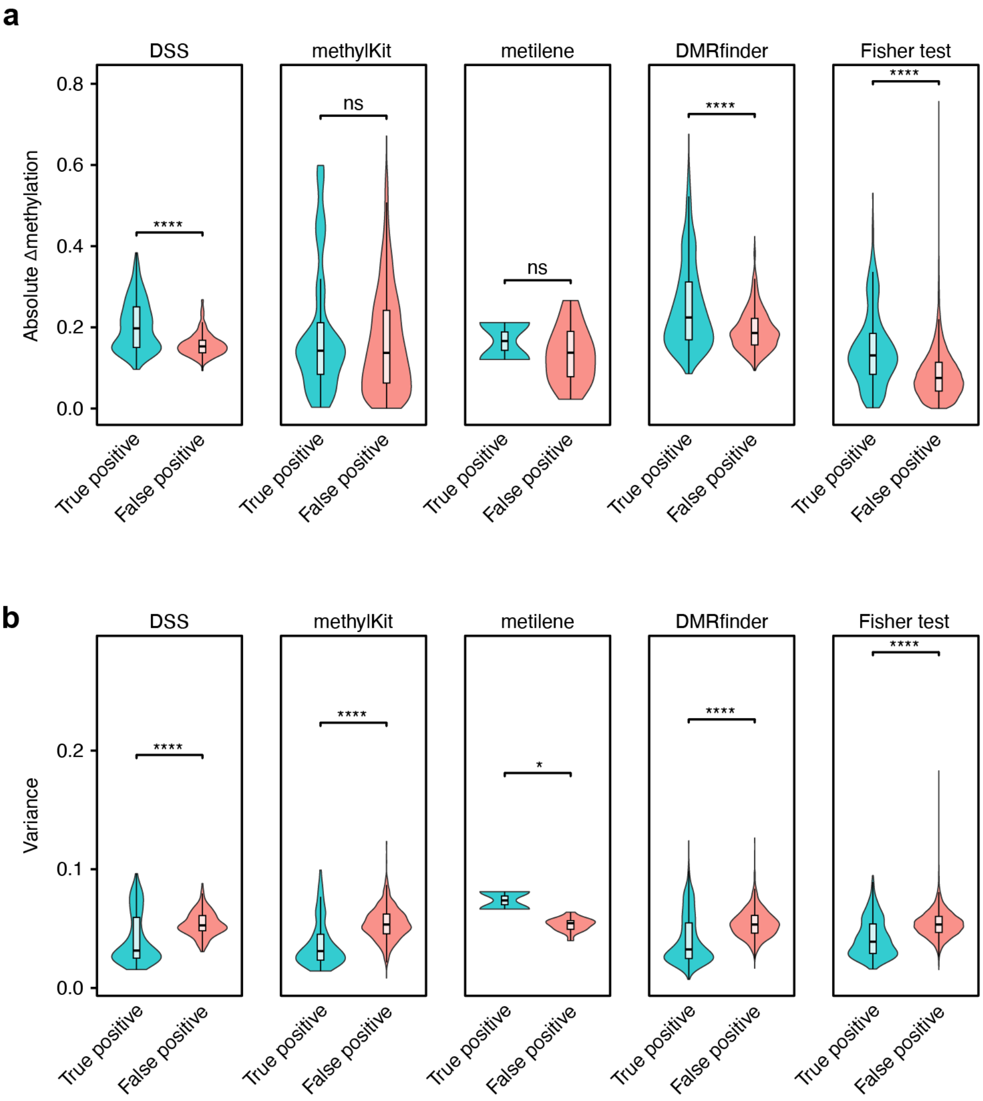


**Fig. S7 Characteristics of true-positive and false-positive DMRs detected by existing methods**
(**a**) Distribution of absolute methylation differences between groups for true-positive and false-positive DMRs detected by existing DMR detection methods. (**b**) Distribution of replicate-level methylation variance among biological replicates for true-positive and false-positive DMRs. Violin plots show the full distributions; boxplots show the median and IQR, with whiskers extending to 1.5 × IQR. The significance of statistical differences between the true-positive and false-positive groups was determined using the Wilcoxon rank-sum test (****$p < 0.0001$; ***$p < 0.001$; **$p < 0.01$; *$p < 0.05$; *n.s.*, not significant).

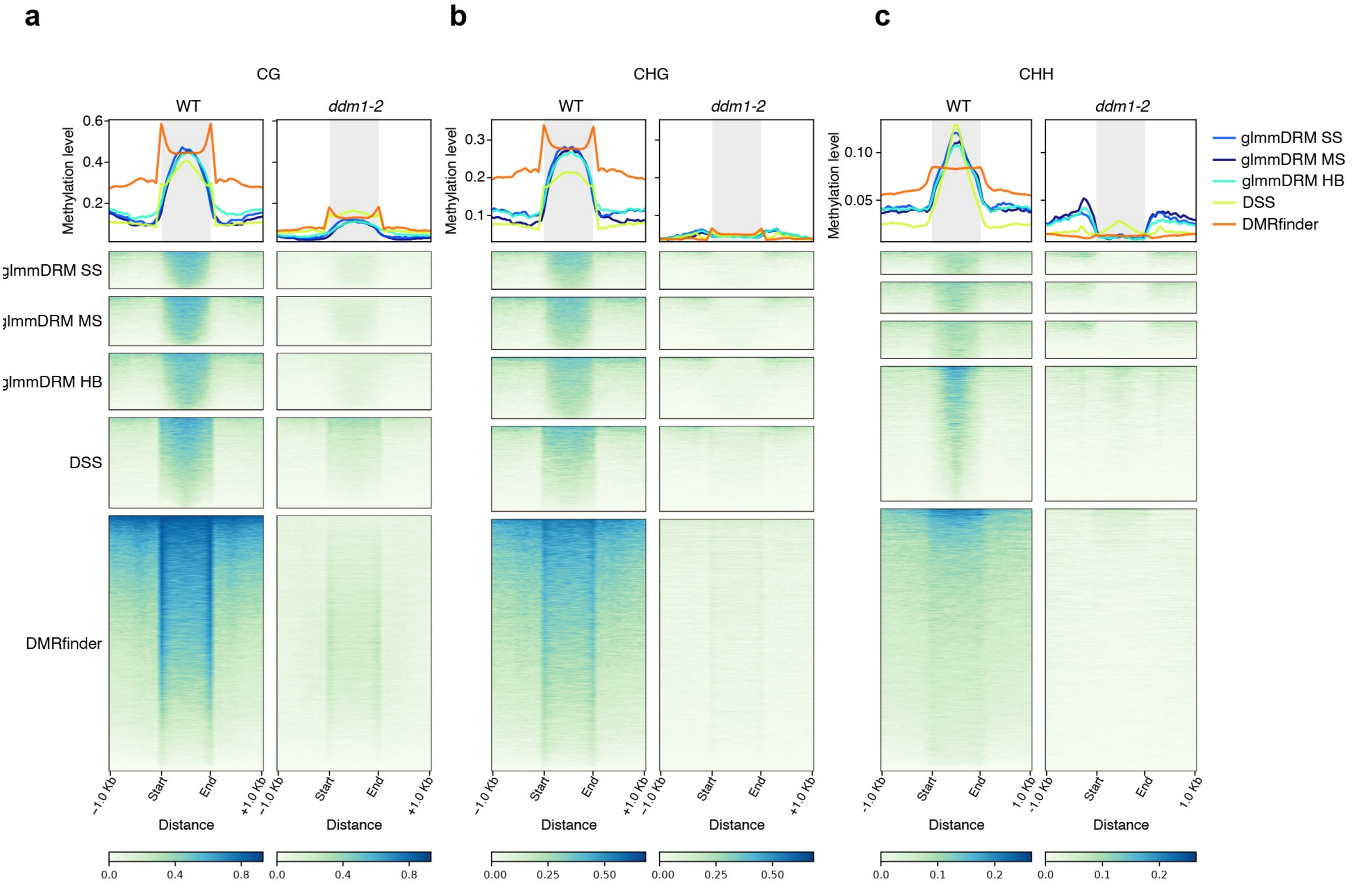


**Fig. S8 Metaplot analysis of methylation levels across detected DMR regions in the methylomes of wild-type Arabidopsis and the *ddm1-2* mutant**

(**a–c**) Metaplots (top panels) and heatmap representation of per-DMR methylation levels (lower panels) of CG (**a**), CHG (**b**) and CHH (**c**) methylation levels across Hypo-DMRs detected by each method in the methylomes of the wild type (WT) or *ddm1-2*. For each methylation context, the DMRs detected by glmmDMR (SS, MS, and HB strategies), DSS, and DMRfinder were used as anchors. The x-axis represents the relative distance between the DMR start and end coordinates, normalized to the same scale, with 1-kb flanking regions on each side. The shaded regions indicate the scaled DMR bodies. In the upper panels, each line represents a different detection method. In the lower panels, rows correspond to individual DMRs and columns correspond to genomic positions along the scaled DMR body. DMRs are ordered by length. The same DMR coordinates were used for WT and *ddm1-2* samples. Color scales indicate methylation levels for each context (CG, 0–0.8; CHG, 0–0.50; CHH, 0–0.2). glmmDMR SS, MS, and HB represent single-seed, multiple-seed, and hybrid strategies, respectively.

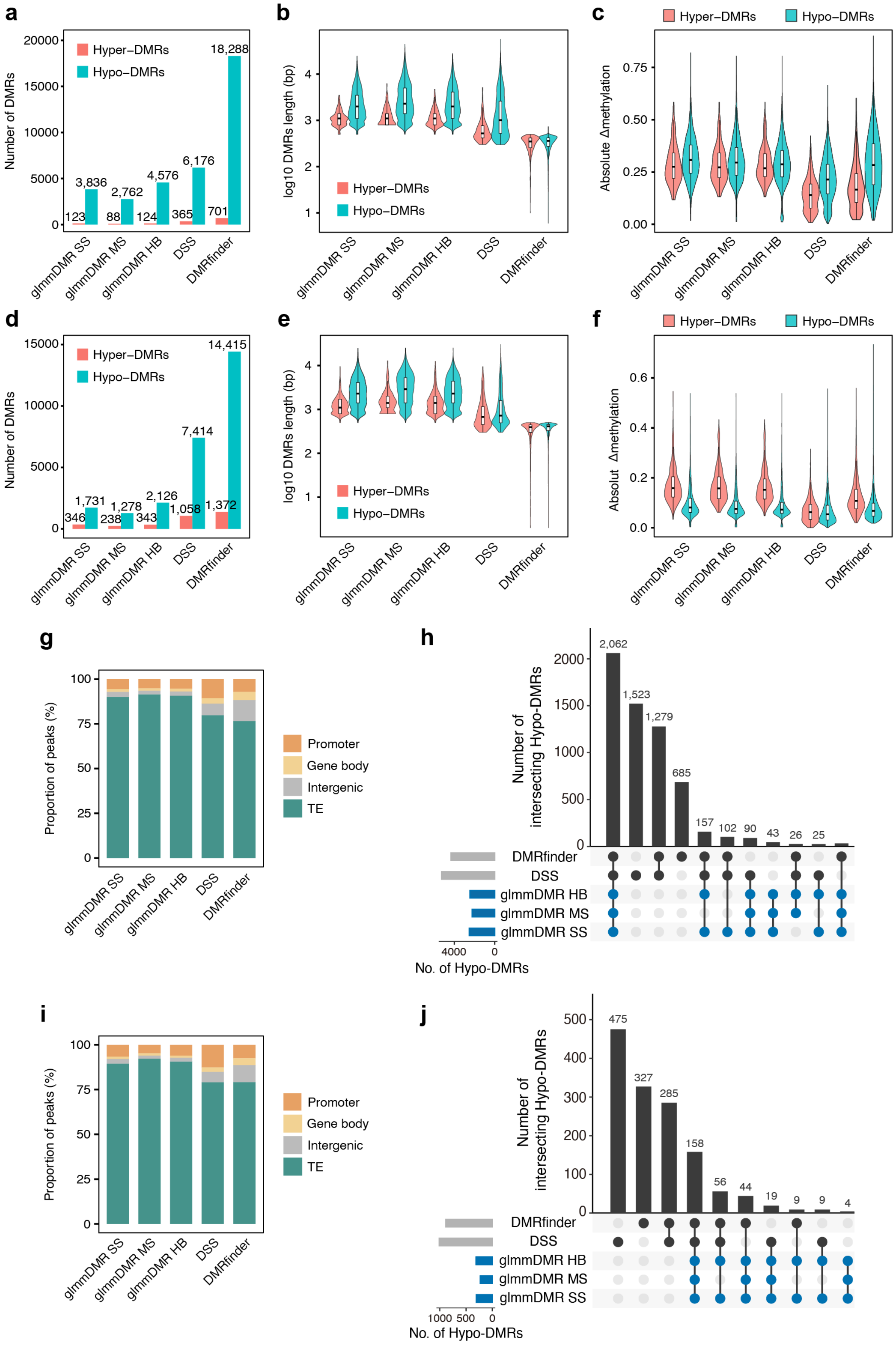
a
Number of DMRs
Hyper−DMRs
Hypo−DMRs
123
3,836
88
2,762
124
4,576
365
6,176
701
18,288
glmmDMR SS
glmmDMR MS
glmmDMR HB
DSS
DMRfinder
b
log10 DMRs length (bp)
c
Absolute Δmethylation
d
346
1,731
238
1,278
343
2,126
1,058
7,414
1,372
14,415
e
f
g
Proportion of peaks (%)
Promoter
Gene body
Intergenic
TE
h
Number of intersecting Hypo-DMRs
2,062
1,523
1,279
685
157
102
90
43
26
25
No. of Hypo-DMRs
i
j
475
327
285
158
56
44
19
9
9
4

**Fig. S9 Comparison of DMR detection methods across CHG and CHH methylation contexts in the *ddm1-2* mutant**

(**a–c**) DMR detection results in the CHG methylation context. (**a**) Number of hypermethylated DMRs (Hyper-DMRs, salmon) and hypomethylated DMRs (Hypo-DMRs, teal) detected by each method. (**b**) Distribution of DMR length (log10[bp]) for Hyper-DMRs and Hypo-DMRs. Violin plots show the full distribution; boxplots show the median and IQR, with whiskers extending to 1.5× IQR. (c) Distribution of absolute methylation differences between group (|Δmethylation|) for Hyper-DMRs and Hypo-DMRs. glmmDMR SS, MS, and HB represent single-seed, multiple-seed, and hybrid strategies, respectively.

(**d–f**) DMR detection results in the CHH methylation context. (**d**) Number of Hyper-DMRs and Hypo-DMRs detected by each method. (e) Distribution of DMR length. (**f**) Distribution of absolute methylation differences between group within detected DMRs. Violin plots show the full distribution; boxplots show the median and IQR, with whiskers extending to 1.5× IQR. (**g, h**) Genomic annotation and overlap analysis of Hypo-DMRs detected in the CHG context. (**g**) Stacked bars showing the proportions of DMRs that overlap with transposable elements (TEs), gene bodies, promoters, or intergenic regions for each method; as in **Fig. 5d**. (**h**) UpSet plot showing the number of shared Hypo-DMRs among glmmDMR configurations (SS, MS, HB), DSS, and DMRfinder; as in **Fig. 5e**. (**i, j**) Genomic annotation and overlap analysis of hypo-DMRs detected in the CHH context. (**i**) Proportions of DMRs that overlap with TEs, gene bodies, promoters, or intergenic regions. (**j**) UpSet plot showing the numbers of Hypo-DMRs shared among methods.

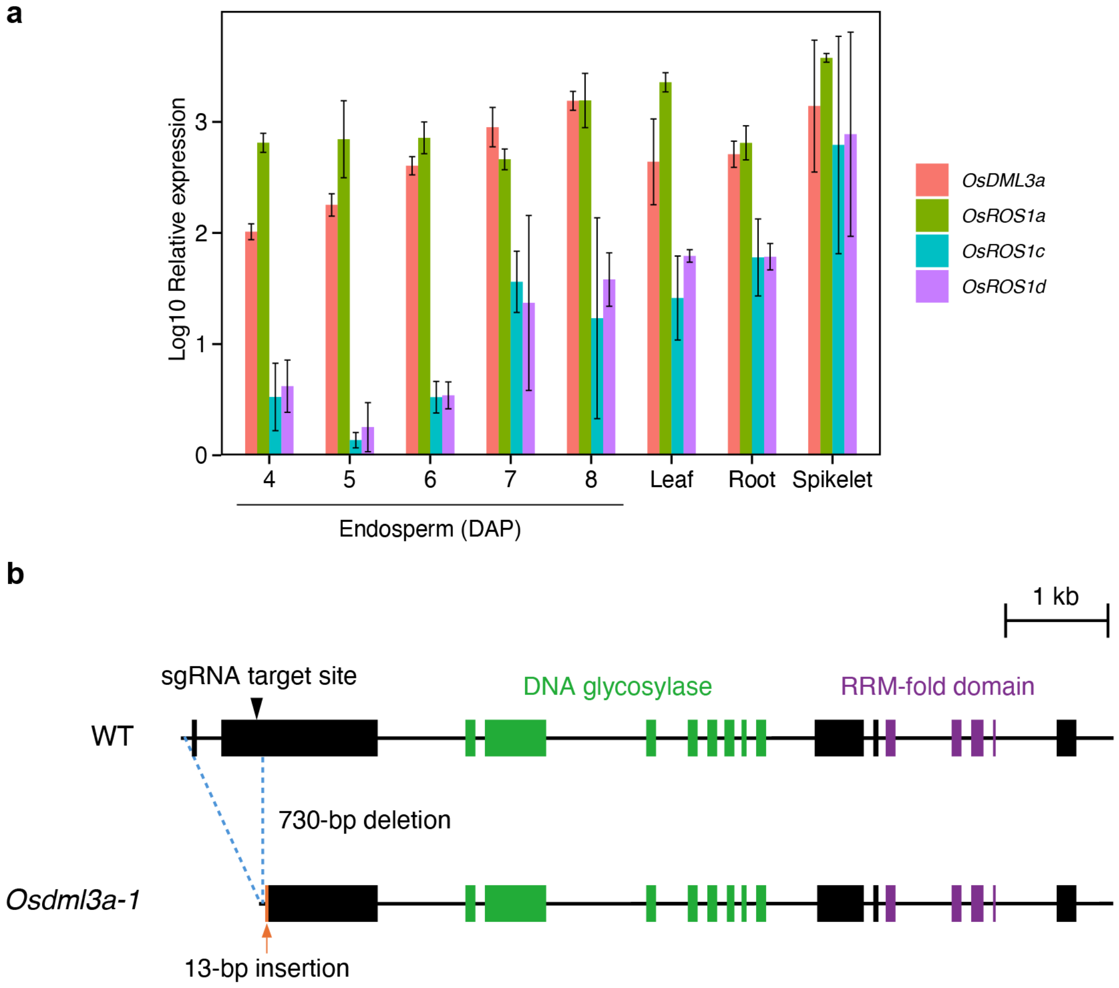


**Fig. S10 Characterization of the *Osdml3a-1* mutant**

(**a**) Spatiotemporal expression patterns of *OsDML3a, OsROS1a, OsROS1c*, and *OsROS1d* determined by RT-qPCR. Expression levels are plotted on a log10 scale and were normalized to those of *UBIQUITIN*. Data are shown as means ± standard deviation SD; ($n = 3$). DAP, days after pollination. (**b**) Diagram of the *OsDML3a* gene structure showing the target site for the single guide RNA (sgRNA) and the genomic alteration in the *Osdml3a-1* mutant. Exons are shown as black boxes and introns as lines. The sgRNA target site is indicated by a black arrowhead. The *Osdml3a-1* mutant carries a 730-bp deletion spanning the upstream region and first exon, together with a 13-bp insertion, resulting in a frameshift predicted to disrupt the OsDML3a open reading frame. The sequences encoding the conserved DNA glycosylase and RRM-fold domains are indicated by green and purple, respectively.

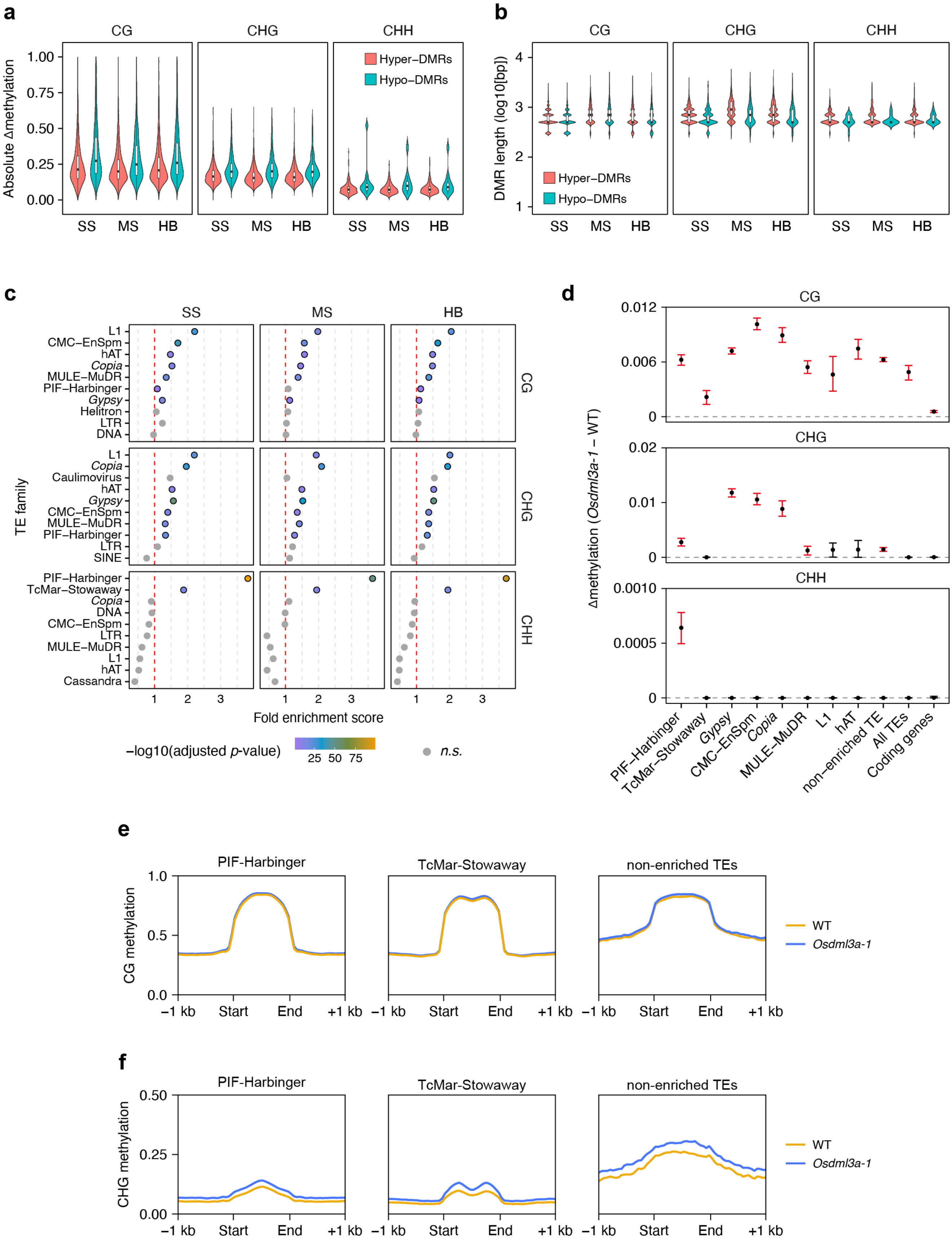
a
CG
CHG
CHH
Hyper–DMRs
Hypo–DMRs
Absolute Δmethylation
1.00
0.75
0.50
0.25
0.00
SS MS HB
b
DMR length (log10[bp])
4
3
2
1
c
SS
MS
HB
TE family
L1
CMC–EnSpm
hAT
Copia
MULE–MuDR
PIF–Harbinger
Gypsy
Helitron
LTR
DNA
Caulimovirus
SINE
TcMar–Stowaway
Cassandra
Fold enrichment score
1 2 3
−log10(adjusted p-value)
25 50 75
n.s.
d
Δmethylation (Osdml3a-1 – WT)
0.012
0.006
0
0.02
0.01
0.0010
0.0005
PIF–Harbinger
TcMar–Stowaway
Gypsy
CMC–EnSpm
Copia
MULE–MuDR
L1
hAT
non–enriched TE
All TEs
Coding genes
e
PIF-Harbinger
TcMar-Stowaway
non-enriched TEs
CG methylation
1.0
0.5
0.0
−1 kb Start End +1 kb
WT
Osdml3a-1
f
CHG methylation
0.50
0.25
0.00

**Fig. S11 DMR characteristics and TE methylation patterns in the endosperm of the *Osdml3a-1* mutant**

(**a**) Distribution of absolute methylation differences between *Osdml3a-1* and wild-type Nipponbare (|Δmethylation|) for Hyper-DMRs (salmon) and Hypo-DMRs (blue) detected by glmmDMR in the CG, CHG, and CHH methylation contexts. Violin plots show full distributions; boxplots show the median and IQR, with whiskers extending to 1.5× IQR. SS, MS, and HB denote single-seed, multiple-seed, and hybrid strategies, respectively. (**b**) Distribution of DMR length (log10[bp]) for Hyper-DMRs and Hypo-DMRs in each methylation context. Boxplots as in (**a**). (**c**) Bubble plot showing the statistical enrichment of TE families among Hyper-DMRs detected by the SS, MS, or HB strategy. Bubbles show fold enrichment, with colors indicating –log10(adjusted *p*-value) calculated by the hypergeometric test. The red dashed vertical lines indicate fold enrichment > 1. For visualization, the top 10 TE families ranked by statistical significance are shown separately for each methylation context and seed strategy. (**d**) Family-wide methylation differences between Osdml3a-1 and wild-type Nipponbare across representative TE families in the CG, CHG and CHH contexts. Points indicate median Δmethylation (*Osdml3a-1* – wild type) calculated using all annotated copies of each TE family, and error bars indicate 95% bootstrap confidence intervals. (**e, f**) Metaplots showing the CG (e) and CHG (f) methylation levels across the bodies of CHH Hyper-DMR enriched (PIF-Harbinger and TcMar-Stowaway) and non-enriched TE families in wild-type (orange) and *Osdml3a-1* (blue) endosperm. The x-axis represents genomic position relative to the TE boundaries. TE bodies were scaled to a common length, with 1-kb flanking regions retained on both sides.
